# Xist-seeded nucleation sites form local concentration gradients of silencing proteins to inactivate the X-chromosome

**DOI:** 10.1101/2020.11.22.393546

**Authors:** Yolanda Markaki, Johnny Gan Chong, Christy Luong, Shawn Y.X. Tan, Yuying Wang, Elsie C. Jacobson, Davide Maestrini, Iris Dror, Bhaven A. Mistry, Johannes Schöneberg, Abhik Banerjee, Mitchell Guttman, Tom Chou, Kathrin Plath

**Affiliations:** Department of Biological Chemistry, University of California Los Angeles, Los Angeles, CA 90095, USA; Departments of Computational Medicine and Mathematics, University of California Los Angeles, Los Angeles, CA 90095, USA; Claremont McKenna College, Claremont, CA, USA; Departments of Pharmacology & Chemistry and Biochemistry, University of California San Diego, San Diego, CA, USA; Division of Biology and Biological Engineering, California Institute of Technology, Pasadena, CA 91125, USA; Keck School of Medicine, University of Southern California, Los Angeles, CA 90089, USA; Molecular Biology Institute, Jonsson Comprehensive Cancer Center, Brain Research Institute, Graduate Program in the Biosciences, Eli and Edythe Broad Center of Regenerative Medicine and Stem Cell Research, David Geffen School of Medicine at the University of California Los Angeles, Los Angeles, CA 90095, USA

**Author notes:** Equal contribution (Johnny Gan Chong, Christy Luong, Shawn Y.X. Tan).

## Abstract

The long non-coding RNA Xist exploits numerous effector proteins to progressively induce gene silencing across the X chromosome and form the inactive X (Xi)-compartment. The mechanism underlying formation of the chromosome-wide Xi-compartment is poorly understood. Here, we find that formation of the Xi-compartment is induced by ∼50 locally confined granules, where two Xist RNA molecules nucleate <u>s</u>upra-<u>m</u>olecular <u>c</u>omplexes (SMCs) of interacting proteins. Xist-SMCs are transient structures that concentrate rapidly recycling proteins in the X by increasing protein binding affinity. We find that gene silencing originates at Xist-SMCs and propagates across the entire chromosome over time, achieved by Polycomb-mediated coalescence of chromatin regions and aggregation, via its intrinsically disordered domains, of the critical silencing factor SPEN. Our results suggest a new model for X chromosome inactivation, in which Xist RNA induces macromolecular crowding of heterochromatinizing proteins near distinct sites which ultimately increases their density throughout the chromosome. This mechanism enables deterministic gene silencing without the need for Xist ribonucleoprotein complex-chromatin interactions at each target gene.

---

Non-coding RNAs are known to seed membraneless bodies in the cytosol and nucleus such as stress granules, the histone locus body, P-bodies, splicing speckles, paraspeckles or Cajal bodies (*1-3*). Their function relies on the recruitment of effector proteins and often on interactions between low-complexity domains that support Liquid-Liquid Phase Separation (LLPS) (*2, 4-10*) A subset of nuclear RNAs, including the long non-coding RNAs (lncRNAs) Xist, Kcnq1ot1, Rox1/2 and Airn, act by regulating chromatin structure and function through the formation of nuclear compartments (*10-21*). The mechanisms underlying compartment formation by this class of RNAs and how they exploit effector proteins to regulate gene expression and chromatin states within the compartment are still poorly understood. Here, we utilize Xist RNA as a model to interrogate these mechanisms. Our work reveals a spatial organization mechanism by which few RNA molecules can regulate a broad nuclear compartment through the recruitment and local concentration of dynamic effector proteins and provides a quantitative framework for studying such compartments.

Xist is transcribed from, coats and silences one of the two X chromosomes during the development of female mammals in a process referred to as X chromosome inactivation (XCI) (*13, 14, 16-18, 21-26*). Xist localization across the X progressively induces changes in gene expression and alters the three-dimensional chromatin structure (*27-29*) through the recruitment of silencing proteins (*30-32*) and the buildup of epigenetic modifications on the chromosome (*33-39*). The prevailing view is that Xist recruits effector proteins to form ribonucleoprotein complexes that spread across the X and silence genes in a stoichiometric manner (*34, 35, 40-42*). However, super-resolution microscopy has revealed that Xist distributes in few diffraction-limited foci on the inactive X chromosome (Xi) (*43, 44*). How a small number of Xist foci recruit interacting proteins to deterministically silence genes across an entire chromosome remains unexplored. Here, we analyzed the distribution and dynamics of Xist and its interacting proteins through quantitative and live-cell super-resolution microscopy, and functionally tested key relationships, to derive fundamental insights into this problem. We performed these analyses during the initiation of XCI in female embryonic stem cells (ESCs), at an early time point when Xist RNA has initially established its territory over the X and silencing initiates at first genes, and at a later timepoint when silencing of most genes has been completed. This critical comparison allowed us to observe changes in Xist and/or protein distributions, mediated by few Xist foci, that lead to robust gene silencing over time.

To explore how Xist RNA orchestrates the progressive formation of the Xi-compartment we first examined when Xist coats the X relative to gene silencing in our ESC to epiblast-like cell differentiation system (*45, 46*) (**Fig. S1A**). RNA FISH showed that Xist demarcated the X-chromosome and formed a large ‘Xist cloud’ by day 2 (D2) whereas silencing of *Atrx* and *Mecp2*, two X-linked genes known to be silenced late during XCI initiation, occurred by day 4 (D4) (**Fig. 1A** and **Fig. S1, B** and **C**). Thus, consistent with prior data (*47*), silencing progresses after Xist initially coats the X chromosome. We therefore chose to examine the X at D2 and D4 of differentiation to understand the mechanisms leading to progressive Xi-compartment formation. We refer to the X state at D2 as the “pre-XCI” state with the “pre-Xi”, and at D4 as the “post-XCI” state with the “Xi”.

**Fig. 1.**
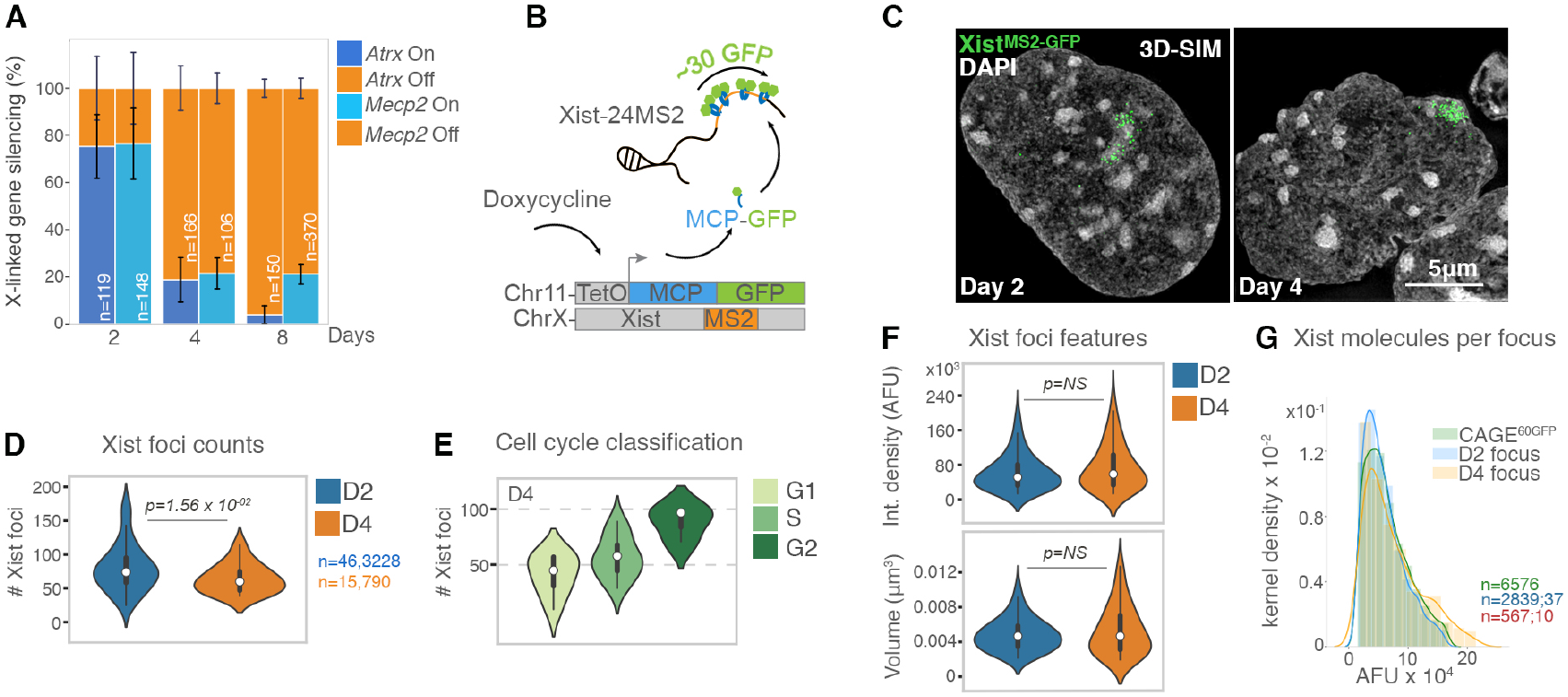
∼50 Xist foci are maintained during progression of gene silencing. **A**, Percentage of cells, from two RNA FISH experiments, with indicated nascent transcription patterns of *Atrx* and *Mecp2* at the indicated day of differentiation. Error bars indicate standard deviation. *n* is the number of cells analyzed. **B**, Illustration of live-cell Xist labelling strategy. **C**, 3D-SIM projections showing Xist^MS2-GFP^ signals (green) and DAPI counterstaining (grey) at the indicated differentiation day. **D**, Violin plots of the 3D-SIM quantification of Xist^MS2-GFP^ foci number at differentiation D2 and D4. Dots denote the median, thick black bars the interquartile range, thin bars show upper/lower values. *n* denotes the number of Xist foci measured (first number) and the number of cells analyzed (second number). Mann-Whitney-Wilcoxon (MWW) *p*-value is given. **E**, As in **D** at differentiation D4 after scoring for cell cycle stages. **F**, Violin plots of the 3D-SIM quantification of Xist^MS2-GFP^ foci showing integrated density of fluorescence (AFU) and volume (μm^3^) of Xist foci at D2 and D4. **G**, Histograms depicting integrated fluorescence of Xist^MS2-GFP^ foci and cages of 60 GFP molecules (cage^60GFP^) (AFU) per pixel kernel density, detected by 3D-SIM.

To determine if the transition from the initial localization of Xist on D2 to the completion of silencing on D4 can be explained by a change in Xist foci number, we used super-resolution three-dimensional Structured Illumination Microscopy (3D-SIM). Previous reports have shown that about 50-200 Xist foci are present in the Xi of female somatic cells (*41, 43*), but whether this number changes during XCI initiation is unknown. To assess Xist foci, we labelled the RNA in living cells by exploiting the MS2 hairpin-MS2 Coat Protein (MCP) interaction (*48*). Briefly, we tagged the endogenous Xist RNA of one of the two X-chromosomes in female mouse ESCs with 24 MS2-repeats and co-expressed MCP-GFP (**Fig. 1B**). MCP-GFP was recruited to Xist (Xist^MS2-GFP^) and the X-linked gene *Atrx* was efficiently silenced (**Fig. S1, D** and **E**), demonstrating the functionality of the Xist^MS2-GFP^ allele. Quantitative 3D-SIM analysis of Xist^MS2-GFP^ shows that the Xist territory consists, on average, of 74 diffraction-limited foci on the pre-Xi and 60 foci on the Xi (**Fig. 1, C** and **D**). These distributions were confirmed by RNA FISH with Xist probes (**Fig. S2, A** and **B, Text S1**). We found that the doubling of the X chromosome with DNA replication in S-phase is accompanied by the doubling of the number of Xist foci, from ∼50 foci in G1 to ∼100 in G2, which was confirmed in somatic cells that maintain the Xi (**Fig. 1E** and **Fig. S2, C** and **D, Text S2**). Thus, the range of Xist foci number in the cell population is largely due to cell cycle differences. Moreover, these findings show that the number of Xist foci is correlated with the length of the X-chromosome. We confirmed this correlation using cell lines that have abnormal Xis of different length (*49*) (**Fig. S2, E** and **F**). Taken together, these data show that transition from the pre-Xi to the Xi is induced by a set number of ∼50 Xist foci.

We next investigated whether the amount of Xist RNA in each focus changes during the transition from pre-to post-XCI. We found that Xist foci are stable assemblies that maintain their integrated fluorescence density and volume during differentiation (**Fig. 1F** and **Fig. S2B**). Consistent with this finding, the Xist locus is constitutively transcribed during differentiation (**Fig. S3, A** and **B**). To estimate the number of Xist molecules in each focus, we utilized a novel fluorescence quantification standard together with quantitative 3D-SIM. Specifically, we transiently expressed nanocages consisting of 60 GFP molecules (cage^60GFP^) (*50*) as internal fluorescence standard in Xist^MS2-GFP^ cells. Integrated density measurements showed that the amount of fluorescence within one Xist^MS2-GFP^ focus on the pre-Xi and the Xi, respectively, corresponds to that of one cage^60GFP^ (**Fig. 1G, Fig. S3, C** and **D**). Fluorescence fluctuation spectroscopy measurements have shown that ∼30 MCP-GFP molecules are bound to 24 copies of the MS2-repeat at a given time (*51*). This number denote that, throughout XCI initiation, each focus contains two molecules of Xist. Thus, only ∼100 Xist molecules orchestrate the initiation of gene silencing across the entire X-chromosome.

The X chromosome contains ∼1000 genes that are subject to silencing (*52*), therefore, the limited number of Xist foci suggests that they should be highly diffusive if they are to directly regulate target genes across the entire chromosome. We therefore investigated the mobility of Xist foci in the pre- and post-XCI states by performing live-cell 3D-SIM of Xist^MS2-GFP^, followed by single-particle tracking of individual foci (**movies S1** and **S2**). This experiment allows for near single-molecule tracking, as each focus contains two Xist transcripts. Unexpectedly, we found that Xist foci exhibited restricted motion and did not undergo fission or fusion (**Fig. 2, A** and **B**). In 90% of cases, the displacement of Xist foci over time was less than 200 nm and their movement was characterized as diffusion in a local confining potential (**Fig. 2, C** to **E** and **Text S3**). The confined motion of Xist foci is highly correlated with the motion of chromatin (*53-57*) both on the pre-Xi and the Xi (**Fig. 2E**). We conclude that Xist foci are tethered to chromosomal locations with high affinity, which constrains each of them locally and limits their movement to that of the Brownian motion of the bound chromatin. Thus, progression of XCI is mediated through ∼50 sites where two Xist molecules are locally confined.

**Fig. 2.**
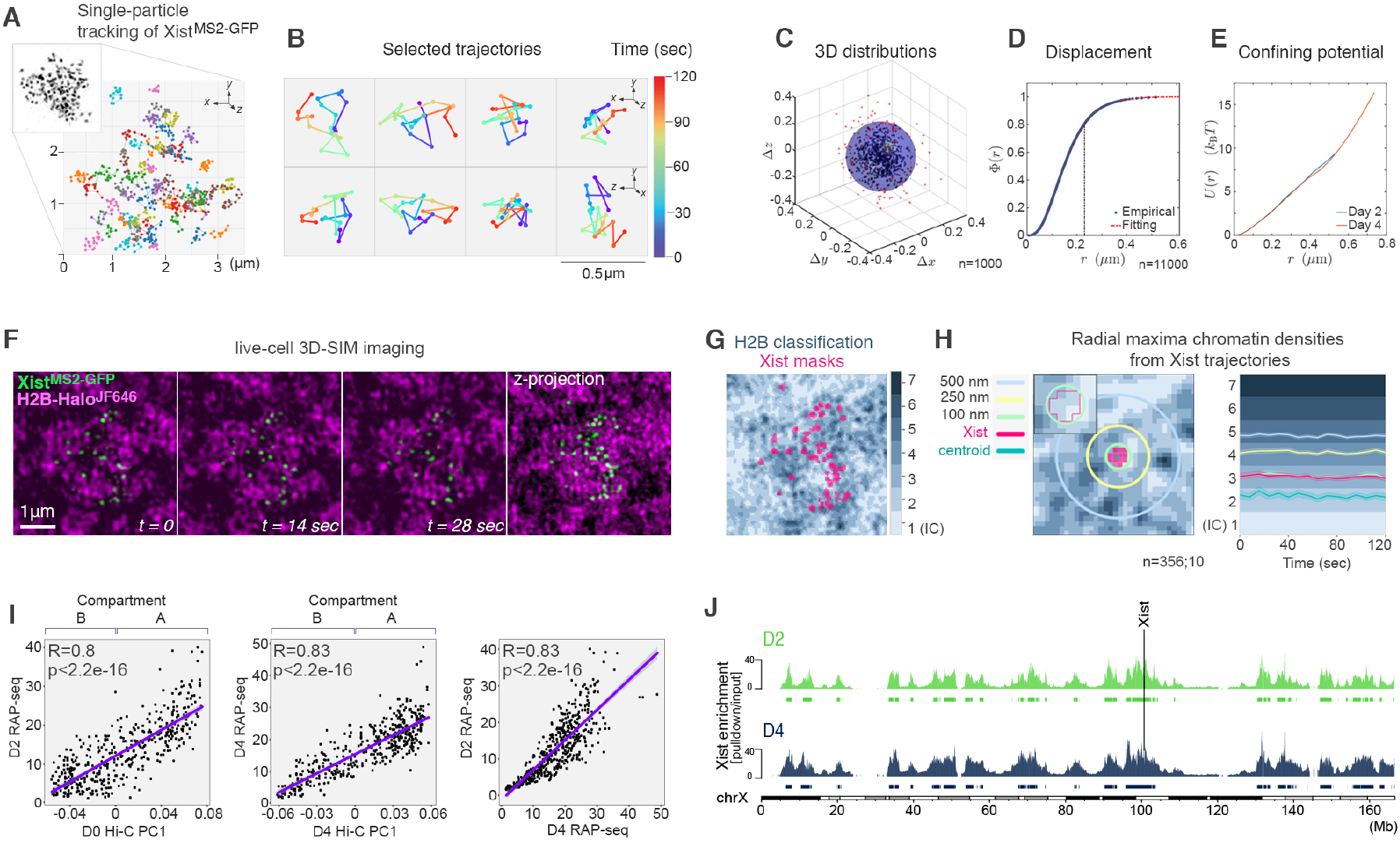
Xist foci are locally confined. **A**, Trajectories of Xist^MS2-GFP^ foci imaged for 2 minutes (5sec/ frame). Inset: Projection of one frame from live-cell 3D-SIM experiment showing an Xist^MS2-GFP^ cluster at D4. **B**, Selected trajectories from **A** showing the displacement of Xist foci over time (color-gradient) in top (xyz, top) and side (zyx, bottom) views. **C**, Coordinates of D4 Xist foci displacements derived from ∼100 trajectories, each centered about their centers of mass. *n* denotes the number of foci analyzed. **D**, Cumulative distribution function *Φ*(*r*) of the number of displacement positions at D4 with distance from origin <*r*. The distance marked with the dashed line at *r*=0.22μm corresponds to the radius of the shaded sphere in **C** where ∼80% of all distances lie (see **Text S3**). *n* denotes the number of foci analyzed from ∼800 trajectories. **E**, Effective spherically symmetric confining potential inferred from the spatial distribution of displacement distances of Xist foci at differentiation D2 and D4. We assume an equilibrium Boltzmann distribution over an effective potential energy well that is a function of *r* (see **Text S3**). **F**, Image sequence from *t*=0 to *t*=28sec and z-projection (from *t*=0) of Xist^MS2-GFP^ (green) and H2B-Halo^JF646^ (magenta), based on live-cell 3D-SIM. Bar: 1μm. **G**, Segmentation of H2B-Halo^JF646^ from live-cell 3D-SIM images into seven density classes and overlay of Xist foci masks. **H**, Left: Schematic for the assessment of the chromatin landscape around one Xist focus. Mask (bright pink) of one Xist focus showing distances (circles) from its centroid (dark green). Inset: magnification showing the outline of the Xist mask. Right: Plot of Xist trajectories showing the average radial maxima of chromatin density reached at each timepoint, where Xist foci never surpass density class 3 and the Xist centroid remains within class 2. Light shaded areas show 95% confidence interval. *n* denotes the number of foci and cells analyzed. **I**, Correlation of Xist enrichment at D2 and D4 of differentiation, based on RAP-seq data, to the first principal component of Hi-C data (A-compartment = positive values, B-compartment = negative values). Far right panel shows the correlation between D2 and D4 Xist RAP-seq data. Pearson correlation r-coefficients and the associated *p*-values are given. **J**, Xist enrichment tracks along the entire X chromosome, defined based on RAP-seq data for Xist over the input, at differentiation D2 (green) and D4 (blue), with peak calls below. The Xist locus is indicated.

To investigate whether the ‘wiggling’ of Xist foci around their centers is confined within a specific chromatin environment, we introduced a histone H2B-Halo transgene into Xist^MS2-GFP^ ESCs and performed live-cell 3D-SIM (**Fig. 2F** and **movie S3**). H2B signals were segmented into seven intensity levels that correspond to chromatin density classes, where class 1 represents DNA-free space (interchromatin channels, IC) and classes 2 to 7 increasing chromatin densities (*43*) (**Fig. 2G**). Xist foci covered predominantly classes 1 to 3 (**Fig. S4A**). Over time, the chromatin densities underlying each focus footprint never surpassed class 3, and the centers of mass (centroids) of Xist foci remained within chromatin class 2 (**Fig. 2H**). These data are consistent with our finding that chromatin density increases in linear increments (**Fig. S4B**). We infer that Xist foci are spatially confined to the periphery of dense chromatin domains, facing the interchromatin channels, and stably maintain their positions relative to chromatin over time. In agreement with these observations, RNA antisense purification (RAP) of Xist from pre- and post-XCI stages followed by DNA sequencing of the associated chromatin (*40*) showed that Xist localizes to gene-rich, open chromatin regions of the A-compartment (*28, 40, 58*) (**Fig. 2I**). The Xist localization patterns are very similar between the pre-Xi and Xi (Pearson’s correlation r=0.83) (**Fig. 2I**). We identified 65 and 63 highly correlated peaks of Xist enrichment on the pre-Xi and Xi, respectively (**Fig. 2J** and **Fig. S4, C** and **D**), similar to the number of Xist foci detected by 3D-SIM. Taken together these findings confirm that the localization of Xist relative to chromatin persists as gene silencing proceeds.

The discovery of only two Xist molecules in ∼50 confined locations indicates that Xist and effector proteins cannot distribute across the entire chromosome space to induce XCI in a stoichiometric manner with gene targets. We reasoned that to effect gene silencing over the entire chromosome, hundreds of Xist-recruited proteins form diffuse clouds localized about Xist foci. These overlapping diffuse clouds can span the entire X and thereby more frequently interact with silencing sites. We addressed this hypothesis by first examining how Xist-interacting proteins accumulate relative to Xist foci, during the initiation of XCI.

We focused on SPEN, PCGF5, CELF1 and CIZ1, four proteins that bind to distinct repeat sequences of Xist RNA and differ in their function in XCI (*59, 60*). SPEN binds the A-repeat sequence of Xist and is the key transcriptional repressor of XCI that activates HDAC3 to induce histone deacetylation and gene silencing (*30, 31, 42, 61-63*). PCGF5 is recruited to the Xi via binding of hnRNP-K to the Xist B/C-repeat sequences (*64*), and is a component of the polycomb complex PRC1 that contributes to the silencing of X-linked genes (*35, 63-67*) and has a critical role in chromatin compaction genome-wide (*68-71*). CELF1 and CIZ1 both bind to the E-repeat sequence of Xist and are critical for restricting the localization of Xist in the X-territory (*72-75*).

We imaged antibody-stained and stably expressed Halo-fusion proteins together with Xist^MS2-GFP^ by 3D-SIM to simultaneously detect Xist and two effector proteins at differentiation D2 and D4 (**Fig. S5A**). Endogenous proteins detected by antibody staining or Halo-fused transgenes displayed similar distributions (**Fig. S5B**). We found that the interrogated proteins formed distinctive assemblies in proximity to Xist foci on the pre-Xi as well as the Xi (**Fig. 3A**). Moreover, protein assemblies appeared larger in the pre-Xi and Xi-territory than in other nuclear accumulations, indicating that Xist induces the *de novo* formation of unique protein complexes. To quantitatively define the protein aggregates induced by Xist, we extracted the spatial coordinates of thousands of diffraction-limited segmented protein foci throughout nuclei (**Fig. S5, C** to **E** and **Table S1**). We measured the nearest-neighbor distances between pairs of different Xist interactors that were either associated with Xist foci or found in the remainder or the nucleus, which includes the active X-chromosome (Xa) (**Fig. 3B**). All investigated pairs of SPEN, CELF1, PCGF5 and CIZ1 foci were, on average, within ∼150-200 nm of each other when associated with Xist foci but separated by >350 nm in the nucleus both pre- and post-XCI (**Fig. 3C**). Therefore, upon distributing across the pre-Xi, foci comprising two Xist molecules immediately recruit arrays of XCI-effector proteins and bring them closer to each other than elsewhere in the nucleus. Hence, large multi-protein assemblies, that are not typically found outside the Xi, form around Xist foci. We refer to these Xist-nucleated proteinaceous nanostructures as Xist-associated supra-molecular complexes (Xist-SMCs).

**Fig. 3.**
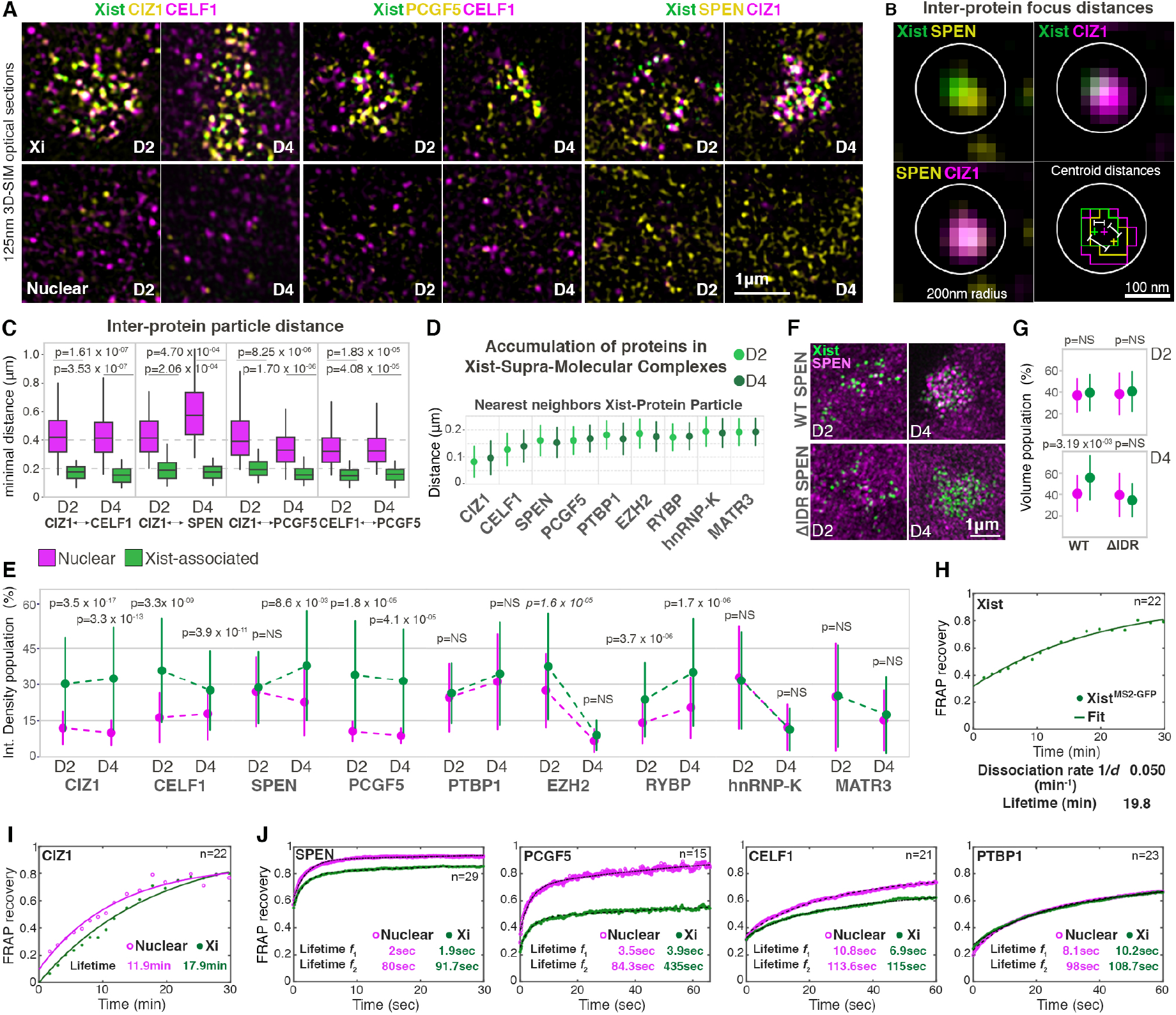
Xist-seeds SMCs of interacting proteins and alters their kinetics. **A**, 125nm 3D-SIM optical sections showing detection of indicated Halo protein-fusions labelled with JF549 (yellow) and immunodected proteins (magenta) in Xist^MS2-GFP^ cells at D2 and D4 of differentiation. Chromatin counterstained with DAPI (grey). Top panels show the Xist-demarcated X-territory (Xi); bottom panels a nuclear region (Nuclear). Note the distinctive enrichment of all pairs of interactors around Xist foci. **B**, Overview of inter-protein particle distance measurements between proteins associated with one Xist focus, based on data in **A**. Overlay: Xist (green), SPEN (yellow) and CIZ1 (magenta). Bottom right panel shows mask outlines after image segmentation, depiction of protein and Xist foci centroids (crosses) and measurement of inter-particle distances as annotated in **C** and **D** are shown. Circle denotes a 200nm radius. **C**, Boxplots showing the distribution of nearest-neighbor distances between the indicated pairs of protein particles in nuclear and Xist-associated fractions (obtained within 250nm from Xist foci centroids) on differentiation D2 and D4. Boxes delineate the upper/lower quartiles, horizontal lines denote the median. *n* denotes the number of protein particles measured; the number of cells analyzed. Mann-Whitney-Wilcoxon (MWW) *p*-value for the comparison between the nuclear and Xist-associated fractions is given. **D**, Point-plots showing the median distance (dot) of the nearest protein particle centroids to Xist particle centroids on D2 and D4. The bars denote the standard deviation. **E**, Point-plots showing the integrated density of fluorescence of indicated protein particles in Xist-associated (green) and nuclear (magenta) fractions, on D2 and D4, from two experiments. Dots denote the median, bars the standard deviation. The medians at D2 and D4 are connected by dotted lines to visualize any changes. Data are normalized to the highest signal observed across the entire population. MWW *p*-values for the comparison between the nuclear and Xist-associated fractions are given. **F**, 3D-SIM projections of the Xist-territory in wild-type (WT) SPEN-Halo ^JF646^ and ΔIDR-SPEN-Halo^JF646^ together with Xist^MS2-GFP^ on D2 and D4. **G**, Particle volume measurements in Xist-associated (green) and nuclear (magenta) fractions, on D2 and D4, comparing ΔIDR-SPEN and WT-SPEN. **H**, Xist^MS2-GFP^ FRAP curve and fitting at D4. Dissociation rate and lifetime inferred from fitting are indicated. **I**, FRAP recovery curve and fitting showing the nuclear and Xist-associated populations of CIZ1-mCherry at D4. Parameters from fitting with a single exponential are given. **J**, FRAP recovery curves and fitting (dashed black lines) showing the nuclear and Xist-associated populations of SPEN-Halo^TMR^, PCGF5-Halo^TMR^, CELF1-mCherry and PTBP1-Halo^TMR^ at differentiation D4. Lifetimes for two subpopulations (*f*_1_, *f*_2_) extracted from bi-exponential fitting are indicated.

To explore whether protein integration into Xist-SMCs is the main mechanism of protein recruitment in the Xi, we probed the distribution of additional XCI effectors (**Fig. S6A**), including the PRC1 component RYBP (*76*); EZH2 (*37, 39, 41, 77*) belonging to the Polycomb repressive complex PRC2; hnRNP-K which binds the B/C-repeats of Xist to recruit PCGF5 (*64*) (reviewed in (*59*)); PTBP1 and MATR3, which function with CELF1 in the maintenance of silencing and Xist localization (*74*). Nearest-neighbor measurements showed a diffraction-limited particle for each of these proteins in a near 1:1 ratio within 200 nm from the center of Xist foci (**Fig. 3D** and **Fig. S6B**). These results corroborate the *de novo* assembly of a multi-protein cloud around Xist foci at the onset of XCI and identify the macromolecular crowding of many direct and indirect Xist interactors in Xist-SMCs, likely involving 100s to 1000s of protein molecules. Notably, our results revealed little differences between Xist-SMCs in the pre-Xi and Xi. Thus, SMCs around Xist foci contain all interrogated proteins before gene silencing completes.

We next investigated whether the pre-Xi to Xi transition is associated with changes in the concentration of proteins within Xist-SMCs compared to the nucleus (**Fig. 3E** and **Fig. S6C, Tables S2** and **S3**). Integrated density and volume particle measurements showed that the levels of CIZ1, CELF1, PCGF5, EZH2 and RYBP are significantly higher in Xist-SMCs than in nuclear foci. For CIZ1, CELF1, PCGF5 levels are stable from D2 to D4, whereas EZH2 and RYBP levels in Xist-SMCs change along with nuclear changes. Conversely, the concentrations of MATR3, PTBP1 and hnRNP-K are similar in Xist-SMCs and nuclear assemblies throughout differentiation, suggesting that they can fulfill their function in XCI at baseline concentration. Overall, these results show that the intense protein accumulations in Xist-SMCs are relatively stable throughout progression of XCI initiation. However, we found that SPEN levels within Xist-SMCs increase from the pre-Xi to the Xi, reaching higher levels than in nuclear assemblies only in the Xi. Thus, the completion of gene silencing is linked to the presence of more SPEN molecules in Xist-SMCs.

We next explored the mechanism that leads to the increased concentration of SPEN in Xist-SMCs with the pre-to post-XCI transition. SPEN contains intrinsically disordered regions (IDRs), which often mediate weak, multivalent interactions (*2, 44, 78-80*). We thus investigated whether the IDRs are required for the progressive accumulation of SPEN. We stably expressed SPEN-Halo with an IDR deletion (ΔIDR SPEN) in Xist^MS2-GFP^ cells and confirmed that it does not interfere with gene silencing (**Fig. 3F** and **Fig. S7, A** and **B**). We found that ΔIDR SPEN and wild type (WT) SPEN accumulated similarly in SMCs on the pre-Xi but, unlike WT SPEN, ΔIDR SPEN levels did not increase on SMCs in the Xi (**Fig. 3, F** and **G, Fig. S7C)**. These results show that the time-dependent increase of SPEN in Xist-SMCs is driven by IDRs and suggest that increased protein aggregation is associated with the completion of gene silencing on the Xi.

Our analyses show that Xist nucleates SMCs, leading to macromolecular crowding of proteins at topologically confined locations. We next explored the kinetic behavior of protein components of Xist-SMCs. We reasoned that the accumulation of proteins in Xist-SMCs should reveal both long-lived binding events, allowing for a topological retention in the SMC, as well as rapidly exchanging constituents, facilitating their access and deposition across the X chromosome. In such a model, transient Xist-SMCs structures would allow SPEN to regulate genes across the entire X chromosome.

To investigate kinetic behavior of protein exchange in the Xi, we stably expressed Halo or mCherry SPEN, PCGF5, CIZ1, CELF1 or PTBP1 fusions, and performed Fluorescence Recovery After Photobleaching (FRAP) over the Xist ^MS2-GFP^ territory or other nuclear regions of the same size. We also examined Xist dynamics for comparison. We observed a slow exchange of photobleached Xist ^MS2-GFP^ (**Fig. S8A**), comparable to previous findings of ectopically expressed Xist (*81*). By fitting measured FRAP curves to a kinetic model with a single-exponential, we inferred a slow dissociation rate (0.05/min) resulting in an average Xist lifetime of ∼20 min, consistent with a single dominant type of high-affinity interaction between Xist and chromatin (**Fig. 3H, Fig. S8B** and **Text S4**). We found that CIZ1 exhibits a ∼18 min recovery time in the Xi, similar to Xist and much longer than that of the other interacting proteins. The lifetime of CIZ1 in the Xi is longer than in other nuclear regions, suggesting that Xist recruitment reinforces CIZ1 binding to chromatin (**Fig. 3I** and **Fig. S8C**). The tight kinetic and spatial (**Fig. 3D** and **movie S4**) relationship between Xist and CIZ1 suggests that CIZ1 and Xist molecules form a stable core of Xist-SMCs.

Kinetic modelling of the SPEN, PCGF5, CELF1 and PTBP1 FRAP curves yielded rapid exchange rates compared to CIZ1 and Xist and two types of binding sites (**Fig. 3J** and **Fig. S9**). Using two-exponential fits we inferred parameters for short-lived (*f*_1_) and long-lived (*f*_2_) bound fractions within and outside of the Xi. For all four proteins rapid binding occurred within seconds and the more stable interactions lasted several minutes (**Fig. 3J**). SPEN is the most dynamic protein, with kinetic rates characteristic of transcription factors (*82, 83*). It is likely that the more dynamic fraction of SPEN (*f*_1_), which has similar characteristics inside and outside the Xi (∼2s), represents fast-exchanging chromatin-engaging molecules that act to control gene expression. Conversely, the longer-lived fraction (*f*_2_) may represent SPEN molecules associated with Xist-SMCs. In line with this hypothesis, FRAP experiments showed a >50% reduction of ΔIDR SPEN of the long-lived Xi binding fraction compared to WT SPEN and a more rapidly dissociating fast population (**Fig. S10** and **Text S5**). Notably, recruitment to the Xi extends the long-lived binding rates and/or increases the fraction of long-lived binding events for the interrogated proteins indicating that the Xi forms a unique nuclear compartment where proteins exhibit distinct kinetic behaviors (**Fig. 3J** and **fig S9, B**). These data are also consistent with their increased accumulation in the Xist territory.

The kinetic assays revealed that proteins with short residence times aggregate around a slowly exchanging Xist-CIZ1 core. These findings denote that Xist-SMCs are rapidly exchanging ‘transient complexes’ that form local, high affinity concentration platforms, thereby increasing the typical residence times of proteins within the Xi. The accumulation in Xist-SMCs primes an increased amount of protein in the Xi, while the rapid binding and dissociation on Xist-SMCs allows proteins, particularly SPEN, to probe and spatially modulate targets further than the locations where two Xist molecules are confined. The presence of a large number of protein molecules in the Xi, without an interaction with Xist, may allow distinct protein species (such as SPEN or PRC1) to explore unique targets and with individual binding rates, which is reflected by the distinct residence time of each tested protein. Such a mechanism is likely thermodynamically favorable and critical for deterministic silencing.

To validate that the formation of Xist-SMCs yields an influx of proteins in the Xi, we next examined the protein population in the Xi but outside Xist-SMCs (referred to as Xi-fraction). We focused on the main two heterochromatinizing proteins SPEN and PCGF5. Using quantitative 3D-SIM, we detected protein assemblies formed in the X-territory, outside Xist-SMCs, which exhibit lower protein concentration than Xist-SMCs but a higher concentration than nuclear assemblies, as assessed by measurements of their integrated fluorescent density and volume (**Fig. S11**). PCGF5 levels in the Xi fraction are significantly higher than within nuclear assemblies on both the pre-Xi and Xi, whereas the level of SPEN in the Xi-fraction increases gradually during this transition. These data suggest that accumulation at Xist-SMCs leads to enrichment of constituent proteins across local neighborhoods in the X, through dynamic feeding from Xist-SMCs.

We next explored the relationship between gradual gene silencing and Xist-SMCs. It is well established that a subset of genes silences soon after Xist coating, whereas other genes, including *Atrx* and *Mecp2*, become silenced later (*29, 34*), yet the mechanism underlying these distinct silencing kinetics is unknown. We examined whether the formation of Xist-SMCs primes early gene silencing events. To test this idea, we investigated whether gene silencing originates at locations proximal to Xist-SMCs by measuring Xist enrichment over gene silencing half-times (*29*). Genes silencing with faster kinetics display more Xist binding on the pre-Xi compared to genes that become silenced later (**fig S12, A** and **B**). These results suggest that rapid silencing kinetics are favorable in proximity to Xist-SMCs. Despite these differences, both fast and slow silencing genes are dependent on SPEN for gene silencing (*30, 31, 42, 61-63*) (**Fig. S12C**). Therefore, we next interrogated whether the ablation of the dynamic association of SPEN with Xist-SMCs interferes with the progression of silencing.

To this end, we exploited an approach (*42*), in which the SPOC domain of SPEN, which is required for transcriptional repression, was tethered to Xist through the BglG/Bgl stem loop (SL) interactions and the endogenous copies of SPEN were depleted (*42, 84*). BglG-BglSL tethering of the SPOC domain to Xist (Xist^SPOC^) is expected to prevent the rapid exchange from SMCs, normally observed for SPEN, and immobilize it primarily at the core of SMCs, where Xist is localized (**Fig. 4A**). If SPEN dynamics control gene inactivation kinetics, then Xist^SPOC^ should result in more efficient silencing of genes that are typically silenced early in XCI than later silencing genes. Indeed, we found that genes normally silenced early during XCI are silenced more effectively by Xist^SPOC^ than genes that are normally silenced late during XCI (**Fig. 4B** and **Fig. S12D**). This result suggests that SPEN-mediated silencing originates at regions that are proximal to Xist-SMCs and expansion of silencing to other target genes occurs progressively and requires the rapid kinetic behavior of the WT SPEN protein. We next addressed the mechanism underlying this expansion.

**Fig. 4.**
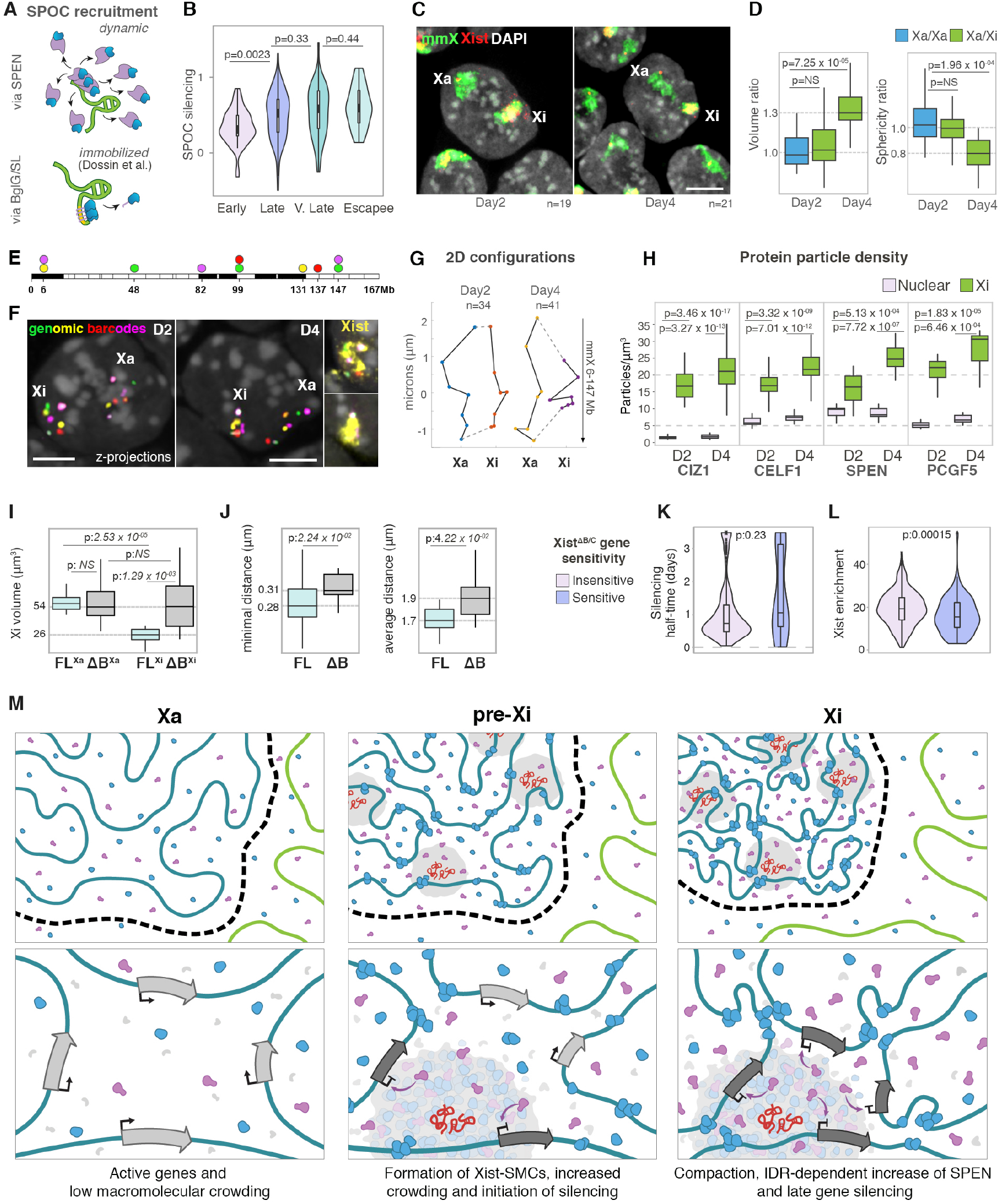
PRC1-mediated compaction is required for late gene silencing by SPEN. **A**, Schematic showing recruitment of the SPEN silencing domain SPOC to Xist under physiological conditions (via the full length SPEN protein) or aptamer tethering (via BglG/SL, Xist^SPOC^). **B**, Violin plots showing the silencing of X-linked genes on Xist^SPOC^ upon depletion of endogenous SPEN. X-linked genes are classified according to their silencing half-times during normal XCI. 0 indicates complete silencing by Xist^SPOC^ and 1 complete lack of silencing. Wilcoxon *p*-values are given. **C**, Projections of confocal stacks of D2 and D4 of differentiation after DNA-RNA FISH using X-chromosome paints (mmX, green) and Xist probes (Xist, red). DAPI counterstaining shown in grey. Bar: 5μm. **D**, Boxplots showing the ratio of volumes (left) and sphericity (right) for Xa/Xa (blue) at day 2 and Xa/Xi (green) at D2 and D4. **E**, Schematic of the spectral barcoding strategy applied to X-chromosome mapping through sequential DNA FISH rounds in **F**. **F**, DNA FISH images of the spectrally barcoded genomic regions on days 2 and 4 of differentiation, as described in **E**. Overlay with Xist RNA FISH signals (far right, yellow) to score for the Xi. Bars: 5μm. **G**, 2D configuration plots of average coordinates of genomic barcodes along the X-chromosome from **F**, extracted with 95% confidence. **H**, Boxplots showing the distribution of the density of indicated protein particles (per μm^3^) in the Xi and in nuclear regions on D2 and D4. **I**, Boxplots showing the volume (μm^3^) of the Xi territory after RNA-DNA FISH with X-chromosome paints and Xist probes in ESCs expressing tet-inducible full-length Xist (FL-Xist) or a deletion-mutant of the B-repeat (ΔB-Xist) after 18hrs of doxycycline induction. **J**, Boxplots showing the minimal (left) and average (right) distance between Xist foci in FL- or ΔB-Xist expressing female cells at D4 of differentiation. **K**, Violin plots showing the silencing half-times (in days) of genes sensitive or insensitive to Xist B/C-repeat deletion. Wilcoxon *p*-value is given. **L**, Violin plots showing Xist RAP-seq enrichment (day 2) of B/C-repeat sensitive/insensitive genes. Wilcoxon *p*-value is given. **M**, Model of XCI. Top panels illustrate part of the X chromosome (blue chromatin) in active (Xa), early silencing (pre-Xi) and inactive (Xi) states and surrounding autosome (green chromatin). Bottom panels show a zoom into the chromatin neighborhood where two Xist molecules (red) nucleate one Xist-SMC. Left: Xa, active genes (light grey arrows). Middle: pre-Xi, macromolecular crowding of Xist-effector proteins (grey, blue, purple proteins) around Xist ‘cores’ leads to the formation of SMCs and initiation of gene silencing (dark grey arrows) in their vicinity. The rapid binding kinetics of proteins creates a local concentration gradient across the entire X-territory and an early Polycomb (blue proteins) deposition across the X. Right: Xi, Polycomb-mediated compaction increases the density of Xist-SMCs, promoting IDR-dependent crowding of SPEN (purple proteins) within Xist-SMCs that leads to an increase in the occupancy of SPEN across the X-territory and completion of gene silencing.

Gradual gene silencing occurs without major changes in Xist-SMC protein levels except for the IDR-dependent increase in SPEN. IDRs are known to adopt more protein-protein interactions upon changes in the local environment (*85-88*), such as an increase in chromatin density that promotes macromolecular crowding (*89, 90*). These observations suggest that the progression of gene silencing beyond targets in the spatial vicinity of Xist-SMCs, as well as the IDR-dependent concentration increase of SPEN, may be mechanistically linked to a conformational change in the X-territory. We therefore investigated whether higher-order chromatin changes occur during the pre-to post-XCI transition.

Measuring the volume and sphericity of the X chromosome upon X-chromosome painting, we found that the conformation of the pre-Xi is similar to that of the active X-chromosome (Xa) (**Fig. 4, C** and **D**). At D4, the Xi reached the distinct compact and spherical organization known for the silent X in somatic cells (*91*) (**Fig. 4, C** and **D**). We extended this result by assessing the conformation of seven unique genomic loci across the X through DNA FISH. We observed a moderate change in the higher-order chromosome configuration of the pre-Xi and a dramatic difference of the Xi compared to the Xa (**Fig. 4, E** to **G** and **Fig. S13, A** to **C**). These changes in chromosomal structure increase the concentration of Xist-SMCs within the X space (**Fig. 4H, Table S4**). Accordingly, minimal distances of Xist foci or protein assemblies in Xist-SMCs are significantly reduced on the Xi compared to the pre-Xi (**Fig. S13D**). Thus, increased chromosome compaction allows more genes to come in closer proximity to the Xist-SMC neighborhood, primes the IDR-dependent SPEN increase in the Xi, and the extension of silencing to more genes over time. However, the mechanisms controlling chromatin compaction on the Xi are not known.

PCGF5 exhibited the highest particle density in the X-territory among all interrogated proteins (**Fig. 4H** and **Fig. S13D**), suggesting that PCGF5-containing Polycomb complexes are significantly more concentrated on the pre-Xi than the other proteins, in agreement with its early occupancy over the chromosome upon induction of Xist (*34, 35*). Given the importance of PRC1 in controlling chromatin compaction and long-range chromatin contacts to regulate gene expression during development (*68-71*), we interrogated whether the extensive concentration of PRC1 on the pre-Xi is critical for inducing the transition of the pre-Xi to a compact Xi.

To test this hypothesis, we functionally perturbed PRC1 recruitment to the X-chromosome by deleting the B-repeat of Xist (ΔB-Xist) in ESCs (*63, 65-67, 92*). Chromosome territory measurements (by X-paints) showed that the volumetric occupancy of an Xi formed by ΔB-Xist is smaller than that formed by the full-length RNA (FL-Xist) and similar to the Xa (**Fig. 4I** and **Fig. S14A**). Consistent with this result, 3D-SIM measurements revealed that the distribution of Xist foci was significantly impaired on a ΔB-Xist Xi compared to a FL-Xist Xi, with larger focus-to-focus distances and an expansion of the Xist-territory, as well as a lack of the characteristic DAPI density of the Xi (**Fig. 4J** and **Fig. S14, B** to **D**). Moreover, the distribution of ΔB-Xist foci on the Xi is similar to that observed on the FL-Xist-coated pre-Xi (**Fig. S13D**). These results uncover a role of the B-repeat, and in turn PRC1, in driving the compaction of the X and the concentration of Xist-SMCs. Consistent with this result, we found that genes that are affected in their silencing by the absence of PRC1, through deletion of the B and C-repeats of the RNA (*65*), are normally more likely to be silenced late during XCI initiation and exhibit lower Xist enrichment (**Fig 4, K** and **L**). These results extend to the architectural protein structural-maintenance of chromosomes hinge domain containing 1 (SMCHD1) that depends on PRC1 for its recruitment to the Xi and controls the compartmentalization of the Xi (*93, 94*) (**Fig S14, E** and **F**). We found that genes with high Xist binding, i.e. close to SMCs, and silenced by Xist^SPOC^ domain are more likely to be SCMHD1-independent, whereas genes controlled by SCMHD1 tend to be inaccessible for silencing by Xist^SPOC^. Thus, SPEN-mediated silencing originates at regions that are proximal to Xist-SMCs and progressive expansion of silencing to other target genes requires the compaction of the X by PRC1 and SCMHD1. Thus, in the absence of PRC1 recruitment to the X, SPEN can still initiate silencing of genes that are exposed to higher SPEN concentrations due to proximity to ∼50 Xist-SMCs, explaining the differential effect on silencing upon SPEN and PRC1 ablation. We conclude that macromolecular crowding, chromatin compaction and silencing by SPEN are interdependent mechanisms of heterochromatin formation.

Taken together, our work reveals a fundamentally new model for how Xist establishes a repressive compartment and orchestrates deterministic transcriptional silencing along the entire X-chromosome (**Fig. 4M**). Our model arises from the key observation that XCI is mediated by a limited number of locally-confined Xist clusters. Through expression, diffusion, sequestration, and degradation (see **Text S6**), Xist becomes localized and tightly bound to chromatin at 50 sites across the X-territory where it strongly interacts with CIZ1 to form the stable core of protein concentration hubs. These Xist/CIZ1 hubs induce macromolecular crowding of multiple XCI effector proteins in their vicinity, prior to chromosome-wide gene silencing and heterochromatinization of the X. In this way, Xist nucleates the formation of SMCs where silencing initiates (**Fig. 4M**, pre-Xi). The high Xist-seeded concentration of PRC1 progressively induces chromatin compaction, altering the local environment of Xist-SMCs and enhancing IDR-dependent protein-protein interactions. Through this process the concentration of SPEN across the Xi also progressively increases, genes move closer to Xist-SMCs and silencing expands across the entire X (**Fig. 4M**, Xi).

The dense seeding of Xist-SMCs across the X-territory and the local binding and unbinding due to transient interactions enriches proteins, and not Xist-ribonucleoprotein complexes, over the X and allows them to spatially probe genomic targets. Consequently, XCI results from locally-confined Xist-mediated macromolecular crowding and supra-molecular aggregation of a large number of protein molecules. Through this process a relatively small number of confined Xist molecules can induce the robust and precise silencing of a much larger number of genes. The dramatic increase in macromolecular concentrations across the X-territory and the sharp change in protein particle density at the boundary of the Xi-territory define the membrane-free chromosome-wide condensate known as the Xi-compartment. The inhomogeneous distribution of Xist-SMCs and altered kinetics of proteins in the Xi, may suggest that Xist induces Polymer-Polymer Phase Separation (PPPS) rather than LLPS in the Xi (*95, 96*).

Finally, our model of how few Xist molecules can establish a chromosome-wide repressive compartment has implications for the regulation of gene expression by other lncRNAs. LncRNAs are typically expressed at low numbers, raising the question of how they can effectively regulate genes. The spatial organization of the X chromosome by few Xist molecules and IDR-based aggregation of protein effectors likely represents a general mechanism through which lncRNAs establish gene-regulatory nuclear compartments. Other lncRNAs have also been found to nucleate spreading of Polycomb complexes (*20*), suggesting that Polycomb-mediated compaction is a common mechanism in the organization of an efficient repressive nuclear compartment.

## Acknowledgements

We thank David Baker (UW) for sharing the ct-60 gene fused to GFP, Joost Gribnau (Oncode Institute) for the MS2-targeting construct, Alexander Shishkin for the SPEN entry clone, Irina Solovei for DNA from flow sorted mouse X-chromosomes, and Yi-Yun Ho for help optimizing the chromosome barcoding experiments. We thank Amy Pandya-Jones for help with the establishment of the ΔB-Xist cell lines and Tsotne Chitiashvili for help with FISH and providing XIST-488 probes. We thank Edith Heard and Francois Dossin for sharing the gene expression profiles after Bgl-GFP-SPOC rescue in SPEN depleted cells. We thank Douglas Black and Emilie Marcus and all members of the Plath laboratory for critical reading of the manuscript and helpful discussions. We also thank the David Geffen School of Medicine (DGSOM) at UCLA, the Jules Stein Eye Institute, David Williams, the Department of Biological Chemistry for supporting the imaging approaches.

## Author Contributions

Y.M. and K.P. conceived the project and Y.M. performed the experiments unless stated otherwise. S.T., C.L., Y.M. and J.C.G. created engineered cell lines and J.C.G., J.W. and Y.M. performed 3D-SIM imaging, data analyses and visualization. K.P. created the ΔB-Xist cell lines. C.L. and Y.M. performed and analyzed FRAP experiments. J.C.G and Y.M. synthesized probes and performed RNA/DNA FISH experiments. T.C. developed all modeling included in this study. E.J. analyzed gene silencing kinetics and correlation of RAP-seq data to Hi-C and expression data. D.M. performed the fitting of FRAP experiments and extracted parameters, and inferred the confining potential of Xist granules, under supervision of T.C. B.M. derived the X-chromosome configurations, which was also overseen by T.C. S.T. performed RAP-seq experiments and I.D. analyzed the data. A.B. cloned the WT- and ΔIDR-SPEN constructs. J.S. wrote the code for subtracting the developmental motion in live-cell 3D-SIM experiments. Y.M., K.P. and T.C. interpreted the data and contributed towards methodology and model creation. K.P. supervised all experimental work. K.P., Y.M., T.C. and M.G. acquired funding to support the project and Y.M., K.P. and T.C. wrote the manuscript with edits from all authors.

## Data and materials availability

All data and material derived from this study are available to researchers upon request.

