## Supplementary information for "Xist-seeded nucleation sites form local concentration gradients of silencing proteins to inactivate the X-chromosome"

#### Materials and Methods

Plasmid construction for engineered cell lines

Plasmids containing the 24xMS2 repeats (#31865) and MS2-Coat-Protein-GFP (MCP-GFP) coding sequence (#27121) were obtained from Addgene. The pBglII5k plasmid was used for targeting the 24xMS2 repeats into Xist (described in (*97*)) and contains homology arms for insertion into exon 7 of Xist, downstream of the E-repeat sequence, and a floxed neomycin resistance cassette. The 24xMS2 repeats were excised from plasmid #31865 by restriction digest with BglII and BamHI and cloned into the pBglII5k plasmid by infusion cloning yielding the pBglII5k-24xMS2 plasmid (which replaces the 16xMS2 repeat array originally contained in the pBglII5k plasmid). The coding region for MCP-GFP was amplified by PCR and introduced under control of a tetracycline-inducible promoter (tetO) into the pBS31 plasmid (pgkATGfrt) (*98*) by infusion cloning yielding pBS31-MCP-GFP. A reverse tetracycline TransActivator (rtTA3) cassette containing the PGK promoter and a BGH polyA element was amplified by PCR from the MXS_PGK::rtTA3-bGHpA plasmid (#62446, Addgene) and introduced into the unique AscI site of pBS31-MCP-GFP, downstream of the tetO-MCP-GFP-polyA insert, by infusion cloning, resulting in the pBS31-MCP-GFP-rtTA3 plasmid. For deletion of the B-repeat of Xist the plasmid was constructed from PCR-amplified 5' and 3' homology regions and a loxP-flanked hygroTK cassette that replaces the B-repeat sequence (chrX: 103480156-103480430).

Genetic engineering strategy for integrating MS2 repeats into the Xist locus in ESCs

The pBglII5k-24xMS2 plasmid was electroporated into the polymorphic *Mus musculus/Mus castaneus* F1 2-1 ES cell line (*99*) after linearization with XhoI. The cell culture was exposed to neomycin selection 36 hours post-electroporation. Colonies were picked and expanded for screening by genotyping PCR and RNA FISH with Xist and MS2 probes. The loxP-flanked neomycin resistance cassette was removed from targeted clones by transient expression of the Cre-recombinase. Subsequently, a FRT-recombination site-containing landing pad was targeted into the *Col1A* locus (on chromosome 11) in F1 2-1$^{24xMS2\text{-}Xist}$ ESCs as described in (*98*). MCP-GFP-rtTA3 expression cassette was then inserted into the FRT site by electroporation of a FlpO-recombinase-encoding plasmid and the pBS31-MCP-GFP-rtTA3 plasmid. The resulting ESC line was denoted as Xist$^{MS2\text{-}GFP}$.

Genetic engineering strategy for deletion of the B-repeat of Xist

F1 2-1 ES cell line (*99*) or male ESCs expressing Xist under a TetO promoter (*40*) were electroporated with linearized plasmid harboring homology arms for targeting into the B-repeat region or XIST and replacing it with a loxP-flanked hygroTK cassette for antibiotic selection. The loxP-flanked hygroTK resistance cassette was removed from targeted clones by transient expression of the Cre-recombinase. Genotyping and confirmation of deletion of the B-repeat on the 129 allele in F1 2-1 ESCs were performed by Southern blotting (not shown).

Establishment of transgenic ESC lines

For the integration of transgenes expressing various mCherry or Halo protein fusions under the control of the endogenous *Rosa26* promoter of $Xist^{MS2\text{-}GFP}$ ESCs, we employed a parent plasmid harboring homology arms for targeting into the *Rosa26* locus and a loxP-flanked puromycin cassette for antibiotic selection (R26 plasmid). A splice-acceptor (SA) and splice-donor (SD) coding sequence synthesized by Genewiz was inserted into the R26 plasmid after MluI/MfeI restriction digest by infusion reaction. The resulting R26-SA/SD plasmid was used as the parent plasmid for insertion of all protein fusions in three-piece infusion reactions. Coding sequence for CIZ1 was amplified from a donor plasmid described in (*74*). Coding sequences for histone H2B and mCherry were amplified from a H2B-mCherry plasmid (Addgene, #20972) and the Halo cDNA was obtained from plasmid Halo-EasyFusion (Addgene, #112852). The coding sequences for the PTBP1, PCGF5, CELF1 were synthesized (Genewiz). To generate the Spen-ΔIDR-Halo plasmid the full-length Spen Entry Clone (Sp22) was modified using Polymerase Incomplete Primer Extension-based mutagenesis with primers designed to delete amino acids 639-3460. Sp22 and the Spen-ΔIDR entry clone, respectively, were inserted into the PyPP-CAG-Halo-V5-IRES-Puro destination vector using Gateway LR Recombination, generating PyPP-CAG-Halo-full-length-Spen-V5 and PyPP-CAG-Halo-Spen-ΔIDR-V5, respectively, both also containing an IRES-puromycin resistance cassette. These plasmids enable constitutive expression of Spen variants with an N-terminal Halo tag and a C-terminal V5 tag and contain a polyoma episomal origin of replication for efficient propagation in mammalian cell culture. All plasmids were verified by restriction digests and sequencing.

Targeting of ESC lines

ESC lines were grown on DR4 feeders. All targetings were performed by electroporation using the GenePulserII (Biorad). Approximately $2x10^7$ cells and 50µg of DNA were resuspended in 400µl PBS in 4mm diameter cuvettes and pulsed twice for 0.2msec at 800V. Antibiotics were added to the growth media 24-36 hours after electroporation. Puromycin was used at 1.5µg/ml, hygromycin at 130µg/ml and G418 at 400µg/ml. The culture medium containing the respective antibiotics was exchanged every 2 to 3 days. Once adequate colony growth was observed (1-2 weeks), 100-200 colonies were picked under a stereoscope, dissociated by trypsinization and seeded in 96-well plate replicates. One replicate plate was used for genomic DNA extraction and subsequent genotyping PCR. All positive clones used in this study were screened to ensure gene silencing by Xist and normal Xist distribution across the X-territory upon induction of differentiation (**Fig. S1, D** and **E**). Additionally, we confirmed that the 24xMS2-repeat unit was introduced into the 129 allele (**Fig. S1E**).

Creation and delivery of the $cage^{60GFP}$ expression plasmid

The gene encoding ct-60 ($cage^{60GFP}$) (*50*) was amplified by PCR. The fragment was introduced under control of the CACGS promoter into the pBS32 plasmid by infusion reaction yielding pBS32-$cage^{60GFP}$ and positive clones were confirmed by restriction digests and sequencing. The pBS32 plasmid was derived from the pBS31 plasmid upon replacement of the tetO promoter with a CAGGS promoter. To visualize both $Xist^{Ms2\text{-}GFP}$ and $cage^{60GFP}$, $Xist^{MS2\text{-}GFP}$ ESCs were differentiated into EpiLCs to induce Xist expression and doxycycline was added to induce MCP-

GFP expression. Expression of the cage$^{60GFP}$ was achieved by transient transfection of the pBS32-cage$^{60GFP}$ plasmid into differentiating cells by Lipofectamine3000 (Thermo Fisher) 24 hours prior to imaging, according to the manufacturer's instructions.

Cell culture

Female mouse F1 2-1 ESCs and its engineered derivatives were grown on 0.5% gelatin-coated flasks seeded with irradiated DR4 feeders (obtained from day 14.5 embryos with appropriate animal protocols in place). Cultures were maintained in mouse ESC medium containing knockout medium DMEM (Life Technologies), 15% FBS (Omega), 2mM L-glutamine (Life Technologies), 1x NEAA (Life Technologies), 0.1mM β-Mercaptoethanol (Sigma), 1x Penicillin/Streptomycin (Life Technologies), and 1000U/mL mouse LIF (homemade) in 5% CO2, 37 °C incubators.

For all differentiation experiments, cells were adjusted for 3 passages to feeder-free conditions in the presence of LIF and two inhibitors, CHIR99021 (3μM) and PD0325901 (0.4μM) (2i+LIF). Epiblast-like (EpiLC) differentiation was performed as described in (*46*). Briefly, cells were maintained for 3 passages in serum-free 2i+LIF N2B27 media containing 1x N2 supplement and 1x B27 supplement (Thermo Fischer), 2mM L-glutamine (Life Technologies), 1x NEAA (Life Technologies), 0.1mM β-Mercaptoethanol (Sigma), 0.5 x Penicillin/Streptomycin (Life Technologies) prior to EpiLC differentiation. To induce differentiation, cells were dissociated and seeded at a density of $2x10^5$ cells/ml in N2B27 media containing 20 ng/ml Activin A (Peprotech) and 12 ng/ml bFGF (Peprotech).

For experiments extending beyond day 4 of differentiation, we applied a protocol previously described in (*100*). Briefly, at day 4 of differentiation, EpiLCs were dissociated with accutase (Life Technologies) and seeded on geltrex-coated coverslips at a density of $5x10^5$ cells/cm$^2$. Cells were then grown in N2B27 media supplemented with EGF and FGF (10 ng/ml each), on geltrex-coated coverslips for 4 more days (d8 of differentiation). At this developmental stage, cells have lost Tsix expression as observed in **Figure S1C**. Media was exchanged daily.

C127 cells were purchased from ATCC and human fibroblasts containing abnormal X-chromosomes (GM3827, GM00735, GM06960, GM07213) were obtained from Coriell. These cell lines were cultured in DMEM (Life Technologies), 15% FBS (Omega), 2mM L-glutamine (Life Technologies) and 1x Penicillin/Streptomycin (Life Technologies).

FISH Probe synthesis

Probes for DNA and RNA FISH experiments and X chromosome paints were labelled by home-made Nick Translation and fluorescent dUTPs as described in (*101*). DNA from flow sorted mouse X-chromosomes was a gift from Irina Solovei. To create mouse Xist probes, we used a full-length mouse Xist cDNA plasmid (p15A-31-17.9kb Xist). Human XIST probes were created from a full-length XIST cDNA construct. For assessing X-linked gene silencing, *Atrx* probes were synthesized using BAC RP23-265D6 and *Mecp2* probes using fosmid WI-894A5. For the chromosome barcoding experiment, we used BACs RP23-53H15, RP23-83J1, RP23-451D5, RP24-81K23, RP24-374B8, RP23-401G5, RP23-104K18. To create an intronic probe against the first intron of

Xist, the corresponding region was amplified from the Xist-encoding BAC RP23-223G18 and was labelled by Nick Translation. RNA or DNA FISH probes were used at a concentration of 0.1μg/cm$^2$. For multispectral chromosome barcoding experiments, individual BACs were labelled separately, pooled in a 1:1 ratio and used at a concentration of 0.2μg/cm$^2$. Nick Translation products were labelled with Atto488-dUTP, Alexa Fluor 568-dUTP, Cy3-dUTP, Cy5-dUTP, Texas Red-dUTP and chromosome paints were labelled with Atto448-dUTPs or Cy3-dUTPs. A summary of the probes used in each experiment can be found in **Table S5**. All BACs and fosmids used in this study were purchased from CHORI-BACPAC.

Halo labelling

For FRAP experiments of Halo-fused proteins, 5μM of TMR Halo ligand (Promega) was added to the culture medium for 30min following a 30min incubation in media without added ligand to wash-off unbound ligand. For fixed and live-cell 3D-SIM imaging, 1μM JF549 or JF646 Halo ligands (Promega) were introduced to the media for 15min, washed-off twice with PBS and exchanged with fresh medium which was incubated for another 15 min. Live-cell imaging or fixation was done as described in the corresponding sections.

Immunofluorescence staining

Immunodetection was performed as described in (*102*). For combined Halo ligand and antibody detection, cells were labelled with the Halo ligands and fixed followed by immunofluorescence staining. For the 4-color 3D-SIM imaging where we detect combinations of proteins together with Xist$^{MS2\text{-}GFP}$ (**Fig. 3A** and **Fig. S5A)** we used CIZ1-Halo and CELF1 antibody staining, SPEN-Halo and CIZ1 antibody staining, PCGF5-Halo and CIZ1/CELF1 antibody staining. Halo transgenes were detected with the Halo ligand JF549 and primary antibodies with secondary antibodies conjugated to AlexaFluor647. In **Figure 3, D** and **E** we used Halo transgenes for detection of CIZ1, WT-/ΔIDR-SPEN, PCGF5, and PTBP1 the Halo ligand JF549 and antibody stainings for RYBP, EZH2, hnRNP-K with secondary antibodies conjugated to CF568 dye. We compared the localization of the CIZ1-Halo fusion protein and the endogenous (antibody-stained) CIZ1 protein and show the same trend (**Fig. S5B**).

RNA/DNA FISH

RNA and DNA FISH experiments were conducted as previously described (*103*). For sequential RNA and DNA FISH experiments with X chromosome paints (mmX paints) and Xist probes, RNA FISH preceded DNA FISH.

For the detection of genomic regions across the X chromosome with spectral barcoding, cells were seeded in ibidi chambers with a gridded bottom and DNA FISH was performed first. 5-color optical z-stacks of 0.35μm were acquired on a confocal Zeiss LSM880 system. Spatial coordinates of the acquired positions were recorded on the ZEN software and saved. Following samples were equilibrated with 50% formamide in 2xSSC pH 7.2 solution for 3 hours at 37$^{o}$C followed by RNA FISH with Cy3-labelled Xist probes. Specimens were returned to the microscope stage and saved spatial coordinates were revisited to acquire the Xist RNA signal and discriminate between the Xi and Xa. Z-stacks from sequential rounds were superimposed using ImageJ/Fiji and alignment of the two sequential rounds was performed with the affine transformation of the StackReg plugin based on the DAPI channel. Although hybridization of RNA usually precedes DNA FISH, we have found that Xist RNA is remarkably stable during the sequential process. Since the sequential hybridization for this experiment was only necessary for the scoring of the Xi, without the need for harsh probe strip-off steps, RNA FISH was performed last.

Cell cycle analysis of Xist foci

To discriminate between different cell cycle stages, we used a combination of EdU pulse labelling, to detect S-phase cells, and anti-histone H3-phospho-Serine10 (Active Motif, #39253), to detect G2/M phase cells, while G1 cells remained marker-free. A 10mM EdU stock solution was diluted 1:1000 in growth media and cells were pulsed for 20 minutes prior to fixation. RNA FISH with Xist RNA probes and detection of EdU by click-iT reaction with CF dye Azide 568 (Biotium, #92082) were combined with immunodetection of phospho-histone H3 Serine 10 as described in (*103*). For the assessment of Xist foci features and number throughout the cell cycle in EpiLCs (at day 4 of differentiation), we used the $Xist^{MS2\text{-}GFP}$ cell line and detected $Xist^{MS2\text{-}GFP}$ signals after addition of doxycycline.

Super-resolution microscopy

3D-Structured Illumination Microscopy (3D-SIM) was performed on a DeltaVision OMX-SR system (Cytiva, Marlborough, MA, USA) equipped with a 60x/1.42 NA Plan Apo oil immersion objective (Olympus, Tokyo, Japan), sCMOS cameras (PCO, Kelheim, Germany) and 405, 488, 642 nm diode lasers and a 568 nm DPSS laser. Image stacks were acquired on the OMX AcquireSR software package 4.4.9934 with a z-steps of 125nm and with 15 raw images per plane (five phases, three angles). Raw data were computationally reconstructed with the soft-WoRx 7.0.0 software package (Cytiva, Marlborough, MA, USA) using a Wiener filter set at 0.001 to 0.002 (up to 0.006 for DAPI) and optical transfer functions (OTFs) measured specifically for each channel using immersion oil with different refractive indices (RIs) as described in (*102, 104*). Images from different channels were registered using alignment parameters obtained from a calibration slide of 100 nm gold grid holes and a second calibration for axial alignment using 100nm diameter Tetraspeck beads (Invitrogen) as described in (*104*).

Live-cell imaging

Wide-field and confocal scanning microscopy (for FRAP experiments) or 3D-SIM live-cell imaging (4D-SIM) were performed at 37°C (for 3D-SIM in conjunction with an objective heater), with 5% $CO_2$, controlled humidity and 10% $O_2$, having equilibrated the system and immersion oils for at least five hours prior to acquisitions. This equilibration was particularly important for obtaining artifact-free 3D-SIM datasets and minimize stage drift. Cells were differentiated in geltrex-coated chambers fitted with a high precision glass (ibidi) with daily exchange of media. To induce MCP-GFP expression, doxycycline was added to the cells two hours prior to acquisitions at a concentration of 1µg/ml. Imaging was performed in media containing no phenol red and supplemented with ProlongLive Antifade reagent (Thermo Fisher). For live-cell 3D-SIM imaging, typically 1µm to 2µm stacks of 125nm z-sections were acquired in 1- or 2-color 3D-SIM imaging to obtain 240-500 raw images per frame in 5-8 second intervals depending on exposure times and z-depth. Photobleaching over time was corrected by using histogram matching on the BleachCorrection plugin in ImageJ/Fiji.

Quantitative 3D-SIM analyses

For image segmentation, 32-bit raw datasets were imported into ImageJ/Fiji (*102*) and converted to 16-bit tiff composite stacks. The segmentation of Xist and protein foci was performed as previously described in (*102*) using the TANGO suite (*105*). Image segmentation pipelines, adjustment of thresholds and creation of seeds were performed in high-throughput batch-processing and without manual intervention. Specifically, raw datasets without filtering or subtraction of signals were imported into the segmentation pipeline. Resulting masks of segmented particles were inspected by overlays over the raw data to ensure that the majority of signals was contained in the area to be analyzed. Nuclear masks were created using the DAPI channel as the segmentation volume. For each channel, a duplicate was generated and filtered with a 3D Gaussian filter with standard deviation of 1 ($\sigma$=1) and a Tophat filter with a radius of two pixels in xy and a one-pixel radius in z. The filtered image was segmented using the 3D Suite's Watershed method. Seed threshold and image threshold for watershed were calculated by equations Mean+StdDev*2*seed multiplier and (Mean+StdDev*2*seed multiplier)/image multiplier (Signal-to-Noise Ratio, SNR) respectively, where seed multiplier and image multiplier were determined and inspected manually to ensure the inclusion of all the regions of interest (ROIs) and the removal of background noise. Object features and distance measurements were performed using the 3D ImageJ Suite's "Measure 3D", "Quantif 3D" and "Distance" option plugins for ImageJ/Fiji.

For the assessment of the cage$^{60GFP}$ versus Xist signals, cells expressing the cage$^{60GFP}$ plasmid were typically imaged in the same Field of View (FOV) as cells with the Xist$^{MS2-GFP}$ signal, allowing us to obtain data that could be directly compared. When cells expressed both entities, since the cages are located in the cytosol, nuclear masks from the DAPI channel were created and Xist$^{MS2-GFP}$ signals were measured inside the masked regions, whereas the signal from the GFP-expressing cages was measured outside the nuclear masks.

To extract global nuclear protein particle features (in and outside the Xi), masks of the protein signals of interest were created by filtering raw data with a 3D Gaussian blur followed by automatic thresholding to include all signals and exclude nucleoli. ROIs within a 4µm radius of Xist centroids were selected for features extraction to limit computation time to ~1 hour per nucleus. Nearest neighbor centroid distances and all distances between ROIs within each channel and across different channels were extracted using the 3D ImageJ Suite for minimal distance and average distance analysis, respectively. Distance averaging was performed in Python. Assignment of Xist-associated signals was based on a proximity threshold to Xist centroids with a radius of 250nm. Signals 500nm away from Xist centroids, resulting in a 'rim' around the Xi due to the scattering of many Xist foci throughout the Xi, were defined as the nuclear fraction. To assign the Xi territory coordinates of protein signals within a 250nm radius of a Xist foci were selected, overlapping (double-called) pixels were removed and multiplied by voxel dimensions. The comparison of protein features, such as integrated density of fluorescence and volume, was performed by measurements acquired in the same laser line (568nm) for all proteins detected either with the Halo ligand JF549 or primary and secondary antibodies conjugated to CF568 dye. For each experiment,

ROIs with integrated density and volume values below the 10th percentile or above the 90th percentile of the dataset were removed as outliers.

Creation of X-chromosome masks and measurements of X-territory volume and sphericity

Confocal optical stacks were imported to ImageJ/Fiji and converted to 16-bit tiffs. Using the Yen method, an automatic threshold was set to create 3D masks for the X chromosome territories. Assignment of the Xi was based on RNA FISH signals from the Xist channel. Masks were imported into 3D Suite and the volume and sphericity measurements of the X chromosomes (Xa and Xi) were extracted. Sphericity is defined as the length of the object over its width, with a maximum value of one. For the creation of mmX masks from 3D-SIM stacks to allocate Xist foci inside and outside the X-territory, a 3D Gaussian blur with standard deviation of 5 ($\sigma$=5) was used to filter the channel with the mmX paint followed by automatic threshold by the Yen method. After creating X-chromosome masks, Xist foci inside the masked region or in the remaining nucleus were analyzed as described in the 'Quantitative 3D-SIM analyses' section.

Analysis of X chromosome conformation from genomic barcoding

Confocal optical stacks were imported into Fiji/ImageJ and smoothed with a 3D Gaussian blur with a standard deviation of 1 ($\sigma$=1) and background removal using the "Subtract Background" plugin with a rolling ball radius of 10 pixels. Xa and Xi (scored by the presence of Xist RNA) were identified and saved as separate stacks. Subsequently, each probe signal centroid was extracted using the "3D Object Counter" plugin. The 3D Object Counter generated a list of coordinates of probe signals for each channel. To assign signals to multi-spectral barcodes consisting of two labels, a nearest neighbor search between the two corresponding channels was applied based on all spatial coordinates in each channel. Once pairs of signals were assigned to the multi-spectral barcodes the coordinates obtained in the shortest wavelength were used. In cases were two adjacent signals were detected per probe, potentially due to the presence of transcripts or DNA replication, only one of the signals was used. The coordinates of individual barcodes for the Xa and Xi at days 2 and 4 of EpiLC differentiation were reoriented in 3D space to compute spatial statistics across all cells. To obtain configurations of chromosomal backbones, for each set of probe coordinates, principal component analysis (PCA) was performed in the x and y axes. The z axis was unused as the segmentation resolution in that axis is significantly lower, contributing to large variations in the z coordinate (*106*) and confounding the reorientation method used which is highly sensitive to anisotropic error. The principle component is assumed to be the "backbone" of the chromosome: the expected orientation of a chromosome if initially stretched out along that component's direction before entropically relaxing into an equilibrium configuration. Each set of probes are rotated in order to align its corresponding principle components with the y-axis and translated such that the probes' centroid is aligned with the coordinate origin. Probes of the same loci were then statistically compared to locate their local spatial centroid and 95% confidence interval for Xa day 2, Xi day 2, Xa day 4, and Xi day 4 separately. Ellipsoids encompassing the 95% confidence interval were plotted around each loci centroid. In order to quantify the relative compaction between Xa and Xi from day 2 to day 4, the pairwise distances of 3D coordinates (x,y,z) between each barcode location were measured and averaged over all cells. Averages of Xa distances were subtracted from those of Xi at day 2 and the same was done for day 4 in order to measure the absolute change between chromosomes. A heatmap of this change was plotted where large negative numbers indicate a higher compaction.

Single-Particle Tracking (SPT) and extraction of trajectories

Individual Xist particles from live-cell 3D-SIM data were extracted by using TrackMate (*107*) an ImageJ plugin. DoG Detector with a 0.2µm diameter was used to define the particles and the Simple LAP Tracker with 0.25 max linking distance, 0.3 gap-closing max distance and 2 gap-closing max frame gap were used to track the particles. Trajectories that were not possible to track for over 10 consecutive frames were not used. Over 850 trajectories from 30 cells were analyzed and approximately 50% of all Xist granules without manual intervention were possible to track per nucleus for an average of two minutes. Data extracted from the software were fed into downstream confinement analyses (see **Text S3**).

Live-cell 3D-SIM image registration: subtracting background developmental motion

Two types of motion are captured simultaneously in live-cell 3D-SIM microscopy: a) the developmental motion of the cell and the nucleus, and b) the individual motion of Xist granules within the nucleus. To specifically extract (b) from live-cell 3D-SIM images, a custom Python/Jupyter tool was created that implemented the following algorithm: 1) A set of Xist granules were tracked using TrackMate. 2) For each timestep $t$, the individual displacement vectors $\boldsymbol{x}_{i,t}$ of each Xist granule were calculated. 3) For each time-step, individual displacement vectors were averaged to obtain $\boldsymbol{X}_t$, an approximation of the developmental motion in that time-step. 4) This developmental motion was then subtracted from each Xist granule's displacement vector to arrive at an approximation to granule $i$ 's motion, $\boldsymbol{x}_{i,t} - \boldsymbol{X}_t$.

Segmentation of H2B density classes and assignment of the maximal radial distances of chromatin density classes to Xist granules

Histone H2B-Halo$^{JF646}$ intensities were extracted on ImageJ/Fiji plugin using the "getValue" macro command, that iterates over every pixel in the image to get the intensity value of each pixel, generating a list of all the pixel intensities and their corresponding coordinates. The list of intensities was imported to Python. Then, seven intensity/density classes of equal variance were determined. The 3D Suite was used to create Xist masks, while Xist trajectories were extracted from TrackMate to obtain spatial coordinates (centroids) from each time point. The matrices were paired within the radius of one pixel and chromatin density classes were measured under the masks. Radial distances were measured at all pixels within the respective 100, 250, 500 nm radius of the Xist centroid and the maximal intensity value within that range was defined. Averaged values were then plotted in a line graph as a function of time. To extract the nearest neighbors in chromatin density maps, neighboring intensities for each H2B pixel were determined as the average intensity of all adjacent pixels and stored in an array. A strip plot was used to plot the averaged intensity values where each value was assigned to one of the 7 classes based on the class of the origin pixel.

Statistical analyses of imaging data and visualization

Data analysis and visualization were performed using Python. All violin plots, boxplots, bar plots and point-plots were generated using Seaborn and Matplotlib. NumPy and SciPy were used for mathematical computation and Pandas for data manipulation and analysis. Unless stated otherwise, all graphs show the median as the central point or the central line, and bars on point plots represent the standard deviation. Point plots of protein integrated intensity and volume in **Figures 3, S7** and **S11** show the percentage of the maximum absolute value in each group. Statistical differences were analyzed by the two-sided Wilcoxon's or Mann-Whitney rank-sum test.

FRAP experiments

FRAP experiments with z-sectioning for $Xist^{MS2\text{-}GFP}$ and CIZ1-mCherry were performed on an LSM880 equipped with an Airyscan on a Plan-Apochromat 63x1.4NA oil immersion objective, an image size of 67.5μm x 67.5μm with a pixel size of 0.085μm. Z-optical stacks of 0.5μm were obtained through a 15μm z-depth. Bleaching was performed in ROIs demarcating the Xist territory or corresponding nuclear (control) regions at full laser power and 4 iterations with a pixel dwell time of 4.04 μsec. The first post-bleach frame was acquired immediately after bleaching. Time series were acquired every 1.3 minutes up to 10 frames and every 2 minutes thereafter for a total of 30 minutes with an Argon ion 488nm laser or a DPSS 561nm laser set to 1% laser power.

Single-plane FRAP experiments for all other proteins were performed on the OMX-SR platform in widefield mode and an image size of 512x512 pixels with a pixel size of 0.08μm. In these experiments we employed transgenic cells lines carrying mCherry-tagged CIZ1 and CELF1 and carrying Halo-tagged SPEN, PCGF5 and PTBP1, respectively (**Figures S8 and S9**). Images were acquired for $Xist^{MS2\text{-}GFP}$ in the 488nm channel (95MHz- 6% amplitude, 20msec) and for all mCherry- or Halo-fused-TMR proteins in the 568nm channel (272MHz, 6% amplitude, 50-100ms exposure). Bleaching in ROIs demarcating the Xist territory or a corresponding nuclear (control) regions was performed by using the 568nm laser line in the Ring-TIRF/PK photokinetics module with a bleach spot of 1μm for one iteration for 0.1 seconds.

FRAP time series from z-stacking were projected and all FRAP data were analyzed as described (*108*). In brief, ~2μm user-defined ROIs identified the bleached region and data intensities were measured through time after normalization for fluorescence decay. To correct for drift, images were registered using the Correct3DDrift plugin. For compiling figures, FRAP time series were bleach-corrected using the BleachCorrect ImageJ/Fiji plugin. FRAP curves for Xist and all proteins were fit to single or double exponential models derived from mass-action kinetics. Squared errors were minimized to obtain best-fit detachment rates, binding site densities, and freely diffusing fractions. Details are given in **Text S4**.

RNA-Antisense purification (RAP-seq)

F1 2-1 female mouse ESCs were seeded on geltrex-coated plates and differentiated for 2 or 4 days. $5x10^6$ cells were collected per conditions after dissociation by accutase and RAP-seq was performed as previously described (*40*). In brief, reads were trimmed using trim_galore (v0.4.1) with default parameters to remove the standard Illumina adaptor sequence. Bowtie2 (v2.2.9) was used to align reads to the mouse genome (mm9) with the default parameters. Reads with mapping quality less than 30 were removed using samtools (v1.2). Picard MarkDuplicates (v2.1.0) was used to remove PCR duplicates. Bedtools intersect (v2.26.0) was used to count reads in sliding windows

(100Kb every 25Kb) along the X chromosome. Xist localization across the X-chromosome was defined by calculating the Xist enrichment scores (pulldown/input) in the sliding windows. Unmappable regions were masked.

Hi-C compartments

To compare Hi-C compartmentalization and Xist enrichment, Hi-C PC1 values for the inactive X were downloaded from GSE99991(*28*). PC1 values at day 0 were correlated with Xist enrichment at day 2 and PC1 values at day 4 with Xist enrichment at day 4. In addition, Xist enrichment at day 2 and day 4 were correlated. Datasets were intersected with plyranges (v1.4.4) (*109*). Pearson's correlation coefficients and p-values (t-test, r≠=/=0) were calculated in R (v3.6.0), and plots were made with ggplot2 (v3.3.2) and ggpubr (v0.4.0).

Xist-tethered SPOC silencing

To identify genes that were repressed by SPOC tethered to Xist in the absence of SPEN, we used allele-specific RNA-seq count data from (*42*). SPOC silencing values for each gene were calculated as described in (*42*). Briefly, we filtered out genes that were skewed or not silenced under control conditions. We then calculated a silencing index under normal conditions ($silencing\_index_{DOX} = 1 - (allelic\_ratio_{DOX}/allelic\_ratio_{control})$ and after degrading SPEN and expressing SPOC tethered to Xist ($silencing\_index_{SPOC} = 1 - (allelic\_ratio_{SPOC}/allelic\_ratio_{control})$. To quantify the silencing defect in cells expressing only Xist$^{SPOC}$, we calculated the SPOC_[silencing_index (SPOC_[silencing_index = 1 – (silencing_indexSPOC + aux/silencing_indexDOX). To determine whether SPOC was more effective at repressing early or late silencing genes, we downloaded the proseq-estimated silencing half-times from (*29*) and categorized genes with a half-time range of (0 - 0.4] days as early silencing genes, (0.4 – 1 days] as late silencing genes (1- 2] days as very late silencing genes, and >2 as escapee genes. Wilcoxon *p*-values were calculated in R. SPEN dependence index and Bgl-GFP [silencing] indices were calculated in the same way.

SMCHD1 sensitivity

To identify genes with a silencing defect as a result of SMCHD1 KO, we downloaded RNA-seq allelic counts from GSE99991 and classified genes as SMCHD1 sensitive or insensitive following the method in (Wang et al). Briefly, %Xi was determined by finding the percentage of reads from the Xi out of the total allele specific reads in each gene. We only included active genes with >13 allele specific reads in all samples, that are normally silenced during X inactivation. SMCHD1 sensitive genes were defined as having %Xi in SMCHD1$^{-/-}$ 3-fold greater than %Xi in WT.

Xist BC-repeat deletion

To identify genes sensitive to Xist BC repeat, we downloaded RNA-seq count data from GSE123743(*65*). We found the log2 fold change of genes with and without dox-induced Xist expression for 2 days using DESeq2 (*110*), for both full length and ΔBC Xist. We then found the difference between the log2 fold changes (Δfc) for full length and ΔBC Xist. We considered genes to be ΔBC sensitive if Δfc was greater than 0.5.

$$\Delta fc = \log_2(\text{Dox 2 days/No dox})\ \Delta BC\ -\ \log_2(\text{Dox 2 days/No dox})\ \text{Full length}$$

We compared the silencing rate of ΔBC sensitive and insensitive genes by intersecting with proseq-identified silencing half-times. We compared the Xist enrichment of ΔBC sensitive and insensitive genes by intersecting with RAP-seq day 2 Xist enrichment using plyranges (*109*). Xist was excluded from these comparisons. Wilcoxon *p*-values were calculated in R.

Antibodies and dilutions

Endogenous CELF1 was detected with monoclonal rabbit anti-CUG-BP1 antibody ab129115 (1/800; Abcam); hnRNP-K with polyclonal rabbit antibody A300-678A (1/800); SPEN (Sharp) with polyclonal rabbit antibody A301-119A (1/1000); MATR3 with polyclonal rabbit antibody IHC-00081 (1/200, all from Bethyl); RYBP (DEDAF) with polyclonal rabbit antibody AB3637 (1/1000; Sigma); Ezh2 with monoclonal rabbit antibody #5246 (1/500; Cell Signaling Technology); CIZ1 with a polyclonal rabbit antibody NB100-74624 (1/800; Novus Biologicals), and histone H3 phospho-Serine 10 with the polyclonal rabbit anti-histone H3-phospho-Serine10 #39253 (1/1000; Active Motif). Two secondary antibodies were used, including high cross-absorbed donkey anti-rabbit IgG CF568 antibody SAB4600076 (1/400; Sigma) and high cross-absorbed goat anti-rabbit IgG Alexa Fluor 647 antibody A21245 (1/400; Life Technologies).

### Supplementary Text

#### Text S1

Explanation of the different output in Xist foci numbers when using Xist RNA probes or MCP-GFP to label Xist

The number of Xist foci is typically lower when detecting the RNA via MCP-GFP compared to detection by RNA FISH using probes that cover the entire spliced transcript. Although Xist foci are diffraction-limited structures, the labelling with FISH probes captures the entire RNA molecule in contrast to MCP-GFP, which detects a region in the last large exon of Xist downstream to the E-repeat sequence. Therefore, RNA FISH-based detection results in slightly larger Xist structures. We suggest that the discrepancy between Xist foci counts is a result of image segmentation retrieving more RNA FISH foci than MCP-GFP foci, presumably due to the ultra-structure of Xist RNA. This difference is particularly evident in S-phase where more RNA molecules are introduced in each focus before complete Xi DNA replication (see also **Text S2**). This observation also suggests that Xist foci have a specific ultrastructure, where the E-repeat of Xist (which is close to the MS2 tag on the linear Xist sequence) is more internally localized within the Xist granule.

#### Text S2

Explanation for the increase in Xist foci and the transient changes in integrated density, volume, and Feret diameter of Xist foci observed during S phase

Quantitative 3D-SIM measurements of Xist foci during the cell cycle show a transient increase in the integrated fluorescence intensity and volume of foci from early- to mid-S phase. This increase possibly arises from the Xist locus replicating prior to complete Xi replication *(111)*. These data suggest that increased levels of Xist production by two Xist transcription loci, prior to full doubling of the Xi DNA, result in the incorporation of additional Xist RNA molecules within existing foci. Moreover, the different average number of Xist foci between cells at d2, d4, d8 of differentiation or in C127 cells, defined across the cell population without discriminating cell cycle stage (**Fig. 1E** and **fig. S2B**), is likely a reflection of different populations of cells across the different cell states. For example, since many more cells are in S-phase at day 4 of differentiation than at day 8, we would expect a higher average count of Xist foci at day 4 than at day 8.

#### Text S3

Xist foci position trajectories and effective confining potentials

Trajectories of individual Xist foci were also tracked. After subtracting the overall developmental motion, three-dimensional coordinates of foci $i$, $(x_i(t_j), y_i(t_j), z_i(t_j))$ were collected at times $t_j$. Since each trajectory of about 10-15 sets of positions of foci are unsynchronized, we assume that all foci are in a local equilibrium position at the start of the measurements. We thus aggregate all position points and assume they are distributed according to some effective potential $U(\mathbf{r})$ where $\mathbf{r}$ is the position of a focus relative to its point of minimum energy, assuming it exists.

Under this local equilibrium approximation, we wish to determine if the effective potential confining the foci changes over time. To further simplify matters, we assume spherical symmetry in $U(\mathbf{r}) = U(r)$ and align the center-of-mass of each trajectory since we assume each trajectory of each focus is sampled from a particle in local thermodynamic equilibrium. After recentering each trajectory, the position vectors $r_i(t_j)\hat{\mathbf{r}}$ can be defined via the magnitude

$$r_i(t_j) = \sqrt{(x_i(t_j) - X_i)^2 + (y_i(t_j) - Y_i)^2 + (z_i(t_j) - Z_i))^2}, \tag{S1}$$

where $(X_i, Y_i, Z_i)$ is the center-of-mass coordinate of trajectory $i$. Taking all positions within a trajectory and across all foci, we have a collection of $N$ points with displacements $r_k$ representing a sample from an underlying distribution $\rho(r)$. Associated with this distribution, we can define an effective energy potential $U(r)$ through $\rho(r) \sim e^{-U(r)}$ (*112*).

To find $\rho(\mathbf{r})$ from our finite position data requires careful regularization or Bayesian approaches (*113,114*). For a crude approximation, we can simply define the cumulative distribution function $\hat{C}(r)$ over the magnitude of displacements $r$ (*112,115*),

$$\hat{C}(r) = \frac{1}{N}\sum_{k=1}^{N}\mathbb{1}(r_k, r), \tag{S2}$$

where $\mathbb{1}(r_k, r) = 1$ if $r_k < r$ and zero otherwise. Mathematically, the cumulative distribution function (CDF) is related to the three-dimensional spherically symmetric density through

$$C(r) = 4\pi \int_0^r \rho(r')r'^2 \mathrm{d}r'. \tag{S3}$$

The inference of $\rho(r)$ and ultimately $U(r)$ from discrete data $\hat{C}(r)$ can be quite ill-conditioned (sensitive to small variations in $\hat{C}(r)$) and some regularization, for example, through through parametric fitting to a smooth function, should be applied. The parametric fitting to $\hat{C}(r)$ is further constrained by the symmetry and analytic properties we impose on the potential $U(r)$. For example, smoothness of $U(r)$ at $r \to 0$ precludes $n \leq 4$ (except for $n = 3$ which leads to a constant in $U(r = 0)$) in the small $r$ property of $C(r) \sim r^n$. Moreover, we need the normalized cumulative $C(r \to \infty) \to 1$. We choose to use a quadratic potential $U \sim r^2$ as a"trial function" and add appropriate terms as corrections. When $U(r) = kr^2$, we find

$$\begin{aligned} C_0(r) &\propto \int_0^r e^{-kr'^2} r'^2 \mathrm{d}r' \propto k^{-3/2}\left[\sqrt{\pi}\mathrm{Erf}(\sqrt{k}r) - 2\sqrt{k}re^{-kr^2}\right] \\ &\approx \frac{r^3}{3k^{3/2}} - \frac{r^5}{5\sqrt{k}} + \frac{r^7\sqrt{k}}{14} - \ldots. \end{aligned} \tag{S4}$$

To this single-parameter ($k$) function, we add the parametric correction function

$$\delta C(r) = \left[a_3 r^3 + \sum_{j=5} a_j r^j\right] e^{-br^2}, \tag{S5}$$

and fit the total function $C(r|\theta) = C_0(r|k) + \delta C(r|a_j, b)$ to the experimental data $\hat{C}(r)$. For our analysis, we used the odd coefficients $j = 5, 7, 9, 11$ and found the best-fit parameters $\theta^* = \{k^*, a_j^*, b^*\}$ to construct the best-fit density

$$\rho(r|\theta^*) \equiv \frac{1}{4\pi r^2}\frac{\mathrm{d}C(r|\theta^*)}{\mathrm{d}r}, \tag{S6}$$

from which we extract the potential $U^*(r) \sim -\log\rho(r|\theta^*)$.

#### Text S4

Mass-action binding and dissociation model of FRAP dynamics

We propose a simple model describing photobleached and fluorescing molecules binding to, and detaching from specific sites such as foci. The fluorescence of Xist in our experiments rely on the fast and strong binding of freely diffusing MCP-GFP to newly produced Xist. An abundance of MCP-GFP ensures that newly produced Xist "instantaneously" fluoresces and that once photobleached, stays photobleached. Consider Xist or one specific Xist-effector protein. Before photobleaching, the concentrations of Xist or Xist-effector proteins are at their specific steady-state values $C_{\mathrm{ss}}$ in the photobleaching region of interest (which typically coincides with the extent of the Xi). We also assume that the binding sites are uniformly randomly distributed in the photobleaching region and that each of them contains many different classes of binding sites, with each class having different molecular attachment and detachment rates. A schematic of the FRAP process is shown in **fig. S8B**. For simplicity, we also assume that molecules bind to these different sites in a parallel manner, not relying on the occupancy of the other types of sites.

In steady state, the overall concentrations of molecules bound to the $i$-type sites are denoted by $C_i(t)$. After photobleaching, the bulk free molecule concentration is replaced by fluorescing protein or Xist that diffuses into the evaluation region. This diffusion process, including binding and unbinding to sites, has been modeled using mass-action diffusion-reaction equations with complex solutions (*116*). Here, we make the approximation that the replacement is fast and that the concentration of fluorescing molecules reaches $C_{\rm ss}^{*} \approx C_{\rm ss}$ very shortly after photobleaching, but before appreciable dissociation has occurred. Thus, at very short times, the fluorescence recovery curves may appear to start at finite levels corresponding to the fluorescence of the free, newly replaced molecules. After this fast replacement, the bulk concentration of fluorescing Xist or protein will be denoted $C_{\rm ss}^{*} = C_{\rm ss}$, where the $*$ indicates a concentration of fluorescing molecules. The time-dependent concentration of fluorescing molecules bound to a type-$i$ binding site is correspondingly denoted by $C_i^*(t)$. The mass-action equations for the photobleached and fluorescing protein concentrations are

$$\begin{aligned}
\frac{\mathrm{d}C_i(t)}{\mathrm{d}t} &= -d_i C_i(t) + k_i(C_i^{(0)} - C_i(t) - C_i^*(t)), \\
\frac{\mathrm{d}C_i^*(t)}{\mathrm{d}t} &= -d_i C_i^* + k_i^*(C_i^{(0)} - C_i(t) - C_i^*(t)),
\end{aligned} \tag{S7}$$

where $C_i^{(0)}$ is the total concentration of available $i$-type sites, $d_i$ are dissociation rates from $i$-type sites, and $k_i \equiv b_i C_{\rm ss}$ and $k_i^* \equiv b_i C_{\rm ss}^*$ are the bulk concentration-weighted attachment rates. The initial condition immediately after photobleaching is $C_i^*(0) = 0$. The initial steady-state fraction of all available sites containing photobleached molecules, $C_i(0) \lesssim C_i^{(0)}$, should be close to the total number of available sites and can be found by considering the steady state before photobleaching (when $C_i^* = 0$) $\mathrm{d}C_i/\mathrm{d}t = -d_i C_i + k_i(C_i^{(0)} - C_i) = 0$, leading to the initial condition $\bar{C}_i = k_i C_i^{(0)}/(k_i + d_i)$.

The concentration $C_{\rm ss}$ of photobleached molecules near granules decreases in time as it diffuses away and degrades. Rather than formulate the full diffusion-degradation equation for $C_{\rm ss}$, as was done in Braga *et al.* (*116*), we will also assume that it is instantaneously replaced and that once a photobleached molecules dissociates, it is lost to the bulk and does not rebind. Thus, we approximate $k_i = b_i C_{\rm ss}(t) \approx 0$ in Eqs. S7 and solve to find $C_i(t) \approx C_i(0)e^{-(k_i+d_i)t}$ and

$$C_i^*(t) = \frac{k_i^* C_i^{(0)}}{k_i^* + d_i}\left(1 - e^{-d_i t}\right). \tag{S8}$$

By including the steady-state bulk fluorescence $C_{\rm ss}^*$ and all possible site types $i$, we can construct the total fractional fluorescence recovery,

$$\begin{aligned}
F(t) &= \frac{C_{\rm ss}^* + \sum_i \left(\frac{k_i^* C_i^{(0)}}{k_i^* + d_i}\right)(1 - e^{-d_i t})}{C_{\rm ss}^* + \sum_i \left(\frac{k_i^* C_i^{(0)}}{k_i^* + d_i}\right)} \\
&= 1 - \frac{\sum_i \bar{C}_i^* e^{-d_i t}}{C_{\rm ss}^* + \sum_j \bar{C}_j^*}, \quad \bar{C}_i^* \equiv \frac{k_i^* C_i^{(0)}}{k_i^* + d_i}, \\
&\equiv 1 - \sum_i f_i e^{-d_i t},
\end{aligned} \tag{S9}$$

where

$$f_i \equiv \frac{\bar{C}_i^*}{C_{\rm ss}^* + \sum_{j=1} \bar{C}_j^*} \tag{S10}$$

is related to the fractional contribution to the fluorescence from molecules that bind to type $i$ sites. These fitting formulae mirror previous standard FRAP models (*116,117,118*).

As a consequence of the initial steady-state concentrations and the neglect of $C_{\rm ss}(t)$, the FRAP recovery dynamics depend only on the detachment rates $d_i$. A single site type thus leads to a simple exponential rise in FRAP recovery. To allow for a biphasic recovery, we also consider a two binding site model $i = 1, 2$. The slower recovery, ($d_2 < d_1$) would describe the dissociation from stronger binding sites in the protein cluster and may appear as immobile fractions, especially if the FRAP experiment does not extend to long times.

We tried fitting all FRAP curves with a two-rate model (using $f_1, f_2, d_1, d_2$ as the four fitting parameters) and found that for Xist and CIZ1 recovery, only one dissociation rate $d_1 \equiv d$ is observable. The lifetimes $\sim 1/d_1$ for these two components are on the order of many minutes.

Since Xist and CIZ1 recovery is not complete at the end of the experiment, any other possible rate $d_2 \ll d_1$. However, the fractions of these slower sites $f_2$ is estimable by the final recovery level at the end of the experiment (~30min). The Xist and CIZ1 recovery curves clearly have three identifiable features, the "initial" recovery at short times, the exponential rise, and the final recovery level. These three features determine the three parameters $1 - f_1 - f_2$, $d_1$, and $1 - f_2$, respectively.

For all other proteins, there is initially a much faster recovery process on the order of seconds and the FRAP curves exhibit two time scales of recovery. In these cases, a biphasic model provides stable best-fits. This indicates that there is a complex distribution of binding sites on the Xist-SMC for each protein and that they either have different binding strengths or dissociate sequentially (*119*). This heterogeneity in binding sites is effectively approximated by two populations.

#### Text S5

Explanation of ΔIDR SPEN kinetics

Next, we explored how the IDRs of SPEN contribute to the kinetic behavior of SPEN in the Xi. We find that deletion of the IDRs shortens the lifetime of SPEN at fast dissociation sites by nearly two-fold. This feature arises in both the nuclear and Xi fractions (**fig. S10C**). Additionally, a smaller fraction of ΔIDR SPEN exists in the long-lived binding state, which also becomes remarkably more stable compared to WT SPEN. These results show that without its IDRs, SPEN does not engage in binding events in the same way and that the IDRs control the kinetic behavior of SPEN in a complex manner. We suggest that multimerization via IDRs stabilizes SPEN at the most rapidly exchanging sites (the $f_1$ fraction), which may extend the dwell time of WT-SPEN at target genes, as described for other IDR-containing transcriptional repressors *(120,121,122)*. Moreover, the much longer dwell time for the long binding events ($f_2$) observed for ΔIDR SPEN suggests that the exchange of SPEN molecules at Xist SMCs may be unfavored in the absence of IDRs, and that the IDR-dependent multimerization in SMCs is required for the dynamic exchange of SPEN at these assemblies *(123)*.

#### Text S6

Expression-diffusion-degradation model for Xist confinement

While our research uncovered mechanisms of Xist and Xist-effector protein accumulation in the X chromosome, the proposed mechanistic picture ultimately relies on the confinement of Xist to the X chromosome throughout the XCI process. Xist has been shown to be constitutively expressed throughout the inactivation process (*124*) (also see **fig. S3A**) and its concentration will continue to increase, unless its degradation is sufficiently active. Here, we provide a qualitative mechanistic argument for Xist confinement. Although the Xist RNA will have many complex interactions within the X, conservation of their overall numbers requires a balanced degradation of Xist to prevent its continual spread out of the X.

First, assume that Xist molecules do not strongly interact with each other or other proteins. The spread of independent Xist molecules can then be described by an effective, simple diffusion process. Along with degradation and a localized source (from the Xist expression site), a rough description of Xist distribution is a mass-action expression-diffusion-degradation model

$$\frac{\partial C(\mathbf{r},t)}{\partial t} = \nabla \cdot (D(\mathbf{r},t)\nabla C(\mathbf{r},t)) - \mu(\mathbf{r},t)C(\mathbf{r},t) + S(\mathbf{r},t), \tag{S11}$$

where $C(\mathbf{r},t)$ is the concentration of Xist at position $\mathbf{r}$ and time $t$, $D(\mathbf{r},t)$ is the effective diffusivity of Xist in the chromatin environment, $\mu(\mathbf{r},t)$ is the Xist degradation rate, and $S(\mathbf{r},t)$ is the source of Xist. A simplifying approximation is to assume spherical symmetry, that $D$ and $\mu$ are constants, and that the source $S(\mathbf{r},t) = S_0\delta(\mathbf{r})$ derives from a "$\delta$-function" source representing a single expression site at position $\mathbf{r} = 0$ within the X chromosome. Here, and in the subsequent FRAP model, we assume that the Xist cloud is quickly formed and consider the 3D steady state solution

$$C_{\rm ss}(r) = \frac{S_0}{r}\exp\left[-\sqrt{\frac{\mu}{D}}r\right]. \tag{S12}$$

This simple diffusion-degradation picture exhibits an Xist concentration profile that is confined within a radius of $L \approx \sqrt{D/\mu}$. Thus, an X-chromosome size of $L \sim 2\mu$m places a constraint on the ratio $D/\mu$.

Xist RNA has high affinity to chromatin through its binding to proteins that contain DNA binding domains (*124*) which can be thought of as a distribution of association sites. Transport across these association sites can thus be described by a small effective diffusivity that has been measured to be $D \approx 0.005 - 0.02\mu\text{m}^2$/s or $D \sim 5 \times 10^{-11} - 2 \times 10^{-10}\text{cm}^2$/s (*116,124,125*). This effective diffusivity is measured in the presence of strong interactions and the real "free" effective diffusion in the "non-strongly-binding" regions of the chromatin are typically larger. If $D \sim 10^{-9}\text{cm}^2$/s, we would require a free-Xist lifetime of $1/\mu \lesssim 1$min in order for it to be confined to $L$.

This simple conservation argument provides a qualitative mechanism for Xist localization. From this starting point, we can consider other processes such as the binding of Xist to itself or to proteins. These nonlinear interactions would generate sharper confinement of the Xist cloud to the X-chromosome and would be needed to explain the asymmetric distribution of Xist about $\mathbf{r} = 0$, which is typically not in at the center of the X, but at its periphery (see **fig. S3A**). These nonlinear interactions also lead to higher order structures such as multimeric foci or supramolecular complexes of RNA and proteins within which Xist may be very stable.

In the FRAP analyses in **Text S4**, free Xist can exchange with those that are bound. We approximated the free Xist (or protein) concentrations to be approximately constant $C_{\rm ss}(\mathbf{r}) \sim C_{\rm ss}$ within their confined region within the X.

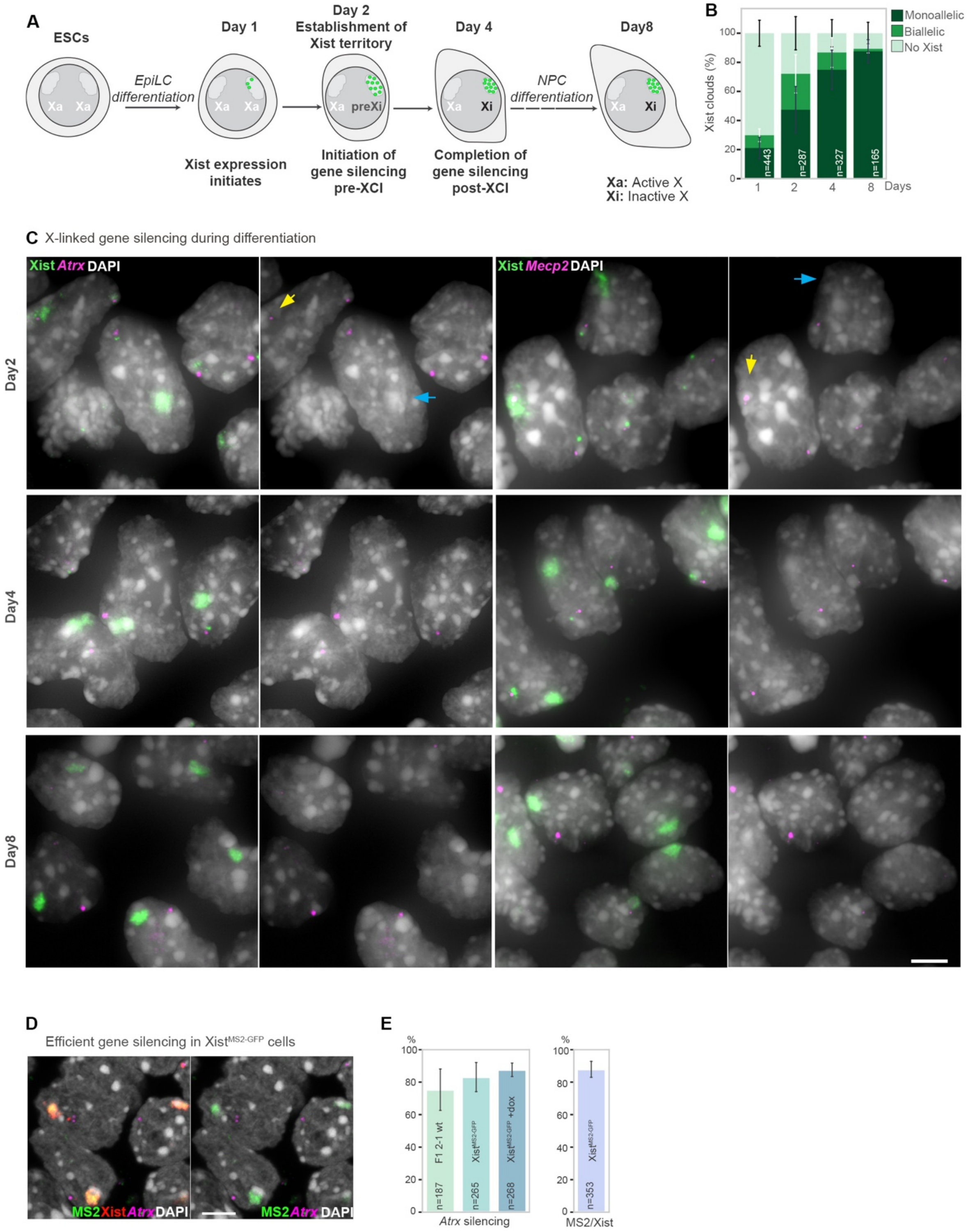
A
ESCs
Xa Xa
EpiLC differentiation
Day 1
Xist expression initiates
Day 2
Establishment of Xist territory
Xa preXi
Initiation of gene silencing pre-XCI
Day 4
Xa Xi
Completion of gene silencing post-XCI
NPC differentiation
Day8
Xa: Active X
Xi: Inactive X
B
Monoallelic
Biallelic
No Xist
Xist clouds (%)
n=443
n=287
n=327
n=165
Days
C X-linked gene silencing during differentiation
Xist Atrx DAPI
Xist Mecp2 DAPI
Day2
Day4
Day8
D
Efficient gene silencing in Xist^MS2-GFP cells
MS2 Xist Atrx DAPI
MS2 Atrx DAPI
E
F1 2-1 wt
Xist^MS2-GFP
Xist^MS2-GFP +dox
n=187
n=265
n=268
Atrx silencing
n=353
MS2/Xist

**Figure S1. Progressive gene silencing after Xist localization on the X chromosome**
**A,** Schematic of ESC differentiation protocol used throughout this study.
**B,** Quantification of the proportion of cells with Xist signals on one or both X-chromosomes at the indicated differentiation day, based on RNA FISH from two experiments. Error bars indicate standard deviation. *n* is the number of cells analyzed.
**C,** Wide-field projections of RNA FISH experiments with Xist RNA (green) and *Atrx* or *Mecp2* (magenta) probes at days 2, 4 and 8 of differentiation, with DAPI counterstaining in grey. The second column of images shows only the DAPI and X-linked transcript channels. Yellow and blue arrows indicate the presence or lack of gene expression, respectively. Bar: 5µm.
**D,** RNA FISH of $Xist^{MS2\text{-}GFP}$ cells at D4 of differentiation with Xist (red), MS2 (green) and *Atrx* (magenta) probes. Chromatin counterstaining by DAPI is shown in grey. The second image lacks the Xist FISH signal. Bar: 5µm.
**E,** Left: Demonstration of effective silencing of *Atrx* in differentiated unmodified (F1 2-1) cells and $Xist^{MS2\text{-}GFP}$ cells at D4 of differentiation, before and after induction of MCP-GFP expression with doxycycline. Right: Quantification of the proportion of $Xist^{MS2\text{-}GFP}$ cells with Xist-MS2 clouds. Error bars indicate the standard deviation.

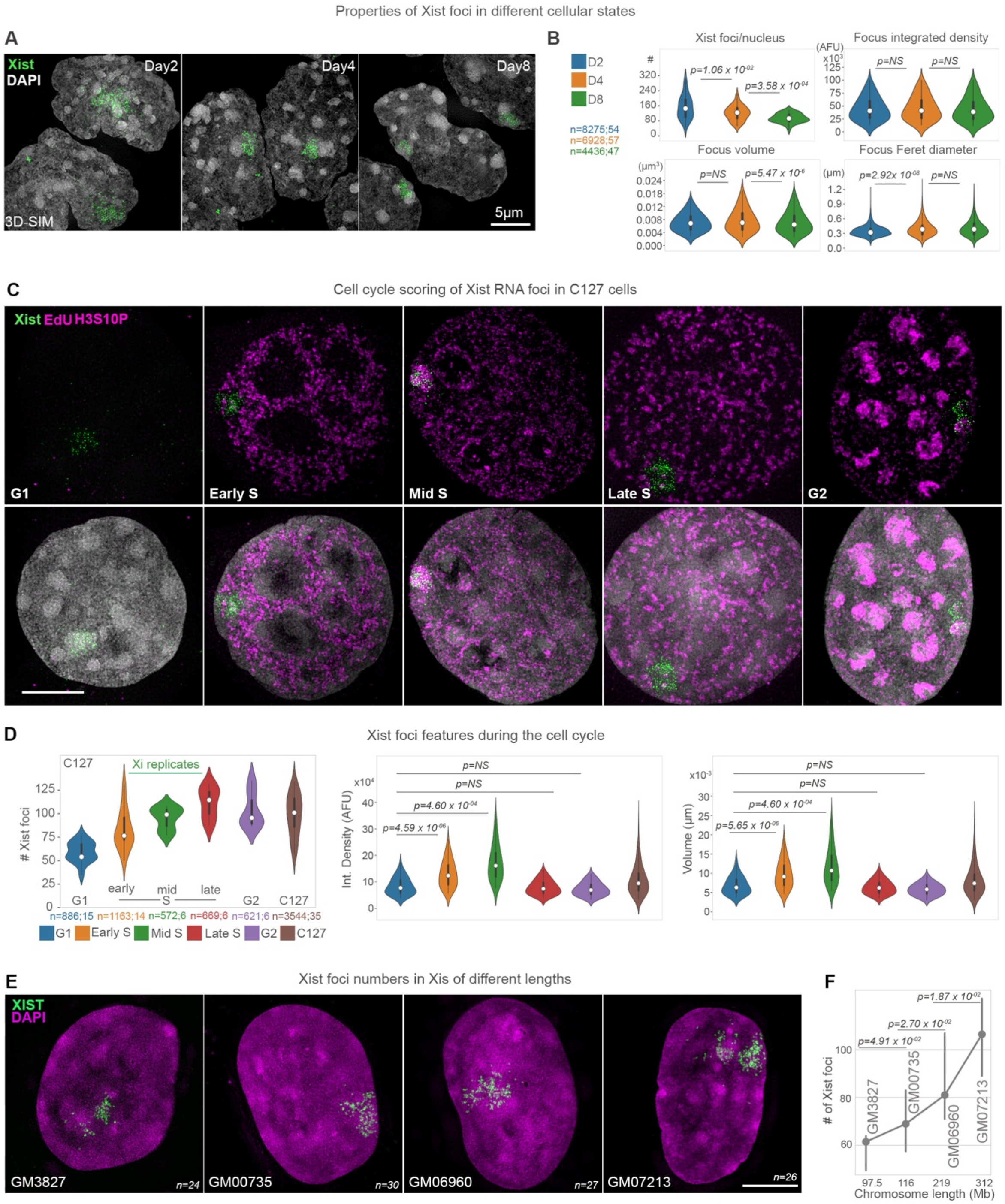


**Figure S2. Features of Xist foci during differentiation, across cell types and X chromosome lengths**

**A,** 3D-SIM projections showing Xist RNA FISH signals (green) and DAPI counterstaining (grey) at the indicated differentiation day. Bar: 5µm.

**B,** Violin plots of the 3D-SIM quantification of Xist foci number (#), integrated density of fluorescence (AFU), volume (µm$^3$), and Feret diameter (µm) at differentiation days 2, 4, and 8 of 2 replicates. White dots denote the median, thick black bars the interquartile range, thin bars show upper/lower values. *n* denotes the number of Xist foci measured; the number of cells analyzed. Mann-Whitney-Wilcoxon (MWW) *p*-values are given by comparing populations indicated by horizontal black lines. NS, Not significant.
**C,** 3D-SIM projections of immuno-RNA FISH experiments detecting Xist, EdU and H3 phospho-serine10 in C127 cells to score for the cell cycle stages G1, early S-phase (Early S), mid S-phase (Mid S), late S-phase (Late S) and G2. DAPI counterstaining in grey is additionally shown in the bottom panels. Stainings for EdU (to detect S-phase cells) and anti-H3 phospho-serine10 antibodies (to detect cells in G2) are detected in the same channel shown in magenta. Bar: 5µm.
**D,** Quantitative features of Xist RNA foci in different cell cycle stages from **C**. The C127-labelled violin plot (brown) depicts the data for the entire cell population. The *p*-value is derived from a MWW test comparing plots as indicated by black horizontal lines. NS, Not Significant.
**E,** 3D-SIM projections of RNA FISH experiments in human cell lines harboring abnormal Xi-chromosomes. XIST RNA probes are shown in green and DAPI counterstaining in magenta. Bar: 5µm. *n* denotes the number of cells used for quantification in **F**.
**F,** Graph showing the average number of XIST granules in each human cell line from **E** (y axis), ordered by the length of the abnormal Xi-chromosome in Mb (x axis). Dots denote the median, bars the 95% confidence interval. *p*-values were derived from a MWW test.

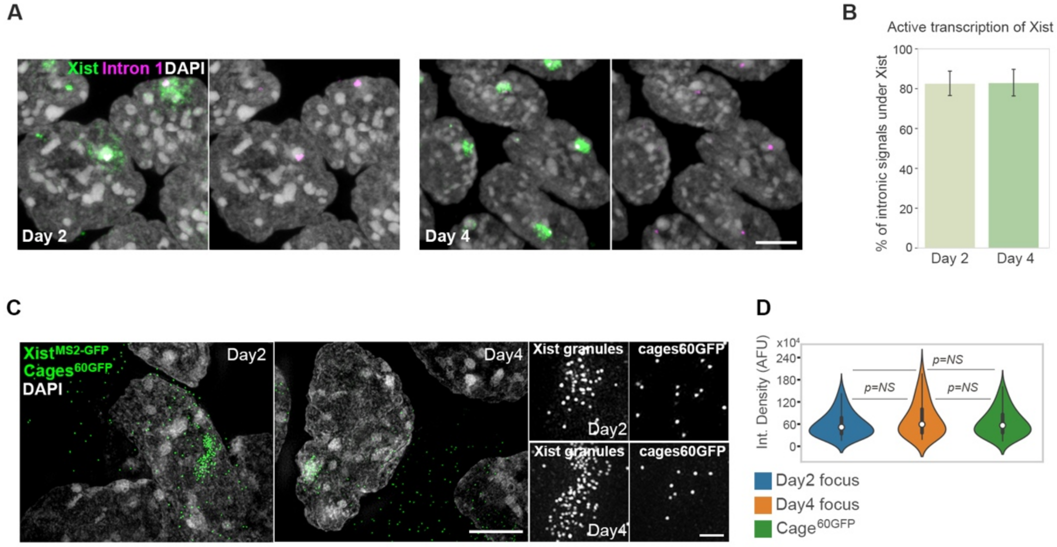


**Figure S3. Stable Xist levels during ESC differentiation**

**A,** RNA FISH with Xist (green) and intron 1 of Xist (magenta) probes to detect nascent transcripts of Xist at differentiation D2 and D4. Chromatin is counterstained with DAPI (grey). The second images show intron 1 and DAPI signals. Bar: 5µm.

**B,** Bar graphs showing the percentages of cells exhibiting a signal for the nascent transcript of Xist under Xist signal-filled nuclear regions (i.e. the pre-Xi/Xi) from experiment in **A**. Error bars denote the standard deviation.

**C,** 3D-SIM projections showing visualization of Xist$^{MS2\text{-}GFP}$ signals (green) at D2 and D4 after addition of 1µg/ml doxycycline for 2hrs to induce MCP-GFP expression. Cells were transfected with plasmids expressing cages$^{60GFP}$ (green) 24hrs prior to fixation. Chromatin is counterstained with DAPI (grey). Bar: 5µm. Bottom panels: magnifications of Xist$^{MS2\text{-}GFP}$ or cages$^{60GFP}$ signals. Bar: 1µm.

**D,** Comparison of integrated density of fluorescence of Xist granules at D2 and D4 and cage$^{60GFP}$ from **C** after image segmentation. The *p*-values from the MWW test were non-significant (NS).

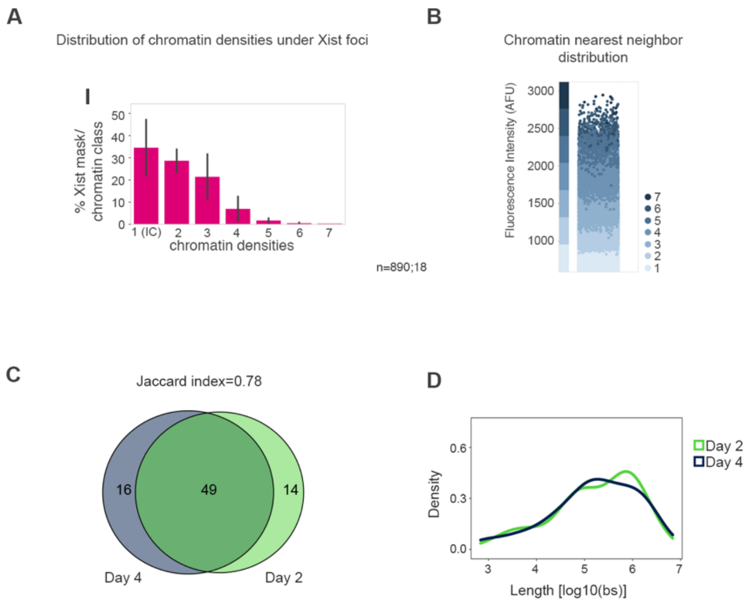


**Figure S4. Persistent localization of Xist in low chromatin density during XCI initiation**

**A,** Quantification of the percentages of chromatin density classes under Xist masks. Error bars denote the standard deviation. *n* is the number of Xist foci analyzed; number of cells.

**B,** Graph indicating the transition between chromatin densities. For each H2B-Halo$^{JF646}$ pixel, the intensity of all neighboring pixels was determined and plotted across the 7 density classes (y axis). The data show that the neighbors (placed in the bin of their associated chromatin density and color-coded by the chromatin class of the pixel of origin) will be found in the same or the directly incrementing density class to the pixel of origin.

**C,** Overlap of Xist RAP-seq peaks across the X-chromosome at D2 and D4 of differentiation.

**D,** Density plot of the length ($log_{10}$(number of bases (bs))) of Xist RAP-seq peaks on the X-chromosome at D2 and D4 of differentiation.

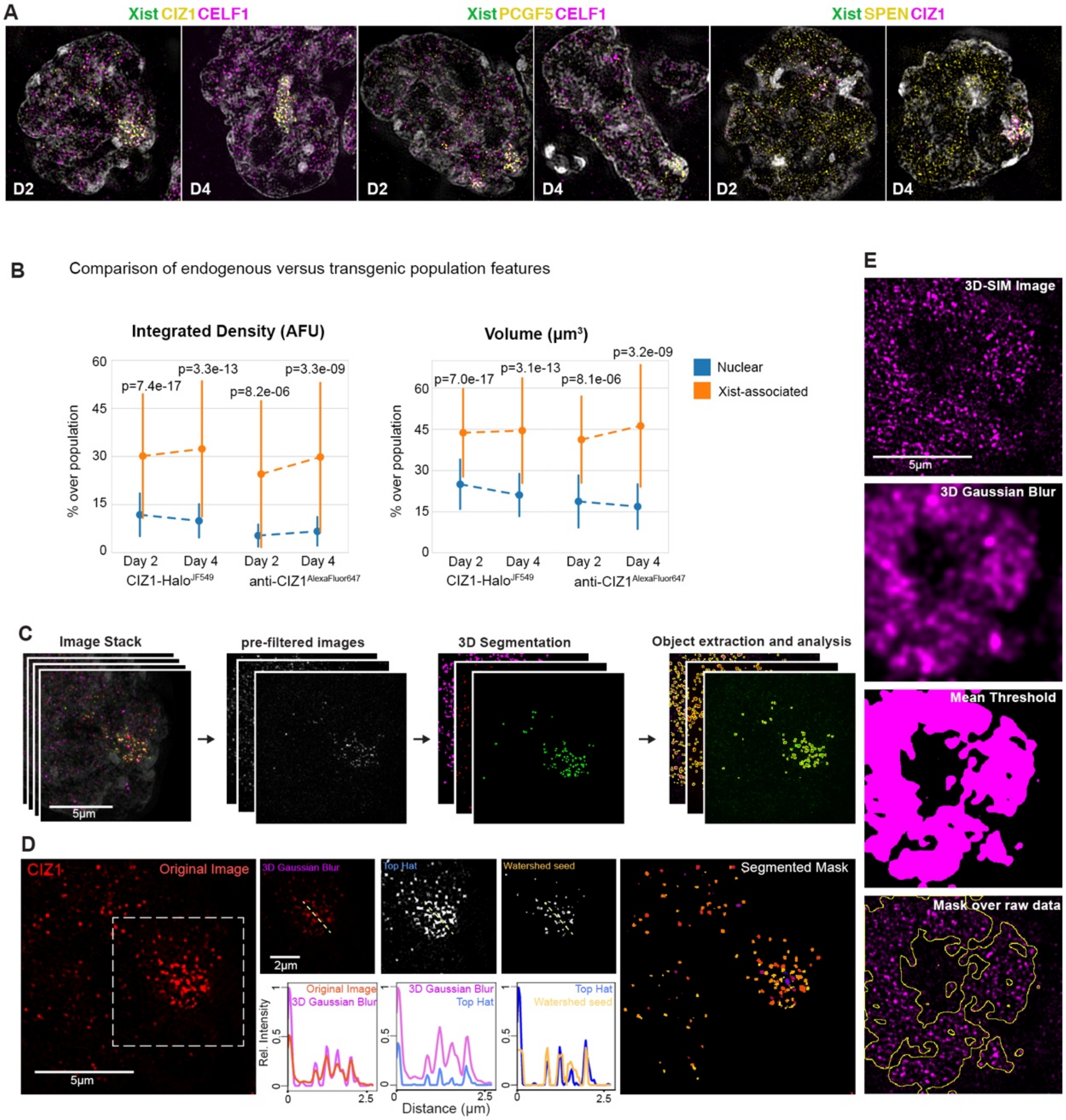


**Figure S5. Quantitative 3D-SIM analysis pipelines**

**A,** 125nm 3D-SIM optical sections showing detection of Halo protein-fusions labelled with JF549 (yellow) and immunodected proteins (magenta) in Xist$^{MS2-GFP}$ cells at day 2 (D2) and 4 (D4) of differentiation. Data are whole nucleus images of insets shown in **Figure 3A**.

**B,** Comparison of the CIZ1-Halo$^{JF549}$ pattern to the localization of the endogenous CIZ1 detected with anti-CIZ1 primary and AlexaFluor647-conjugated secondary antibodies showing the same trend in both endogenous and transgenic protein populations after 3D-SIM quantitative analyses. *p*-values are derived from a MWW test.

**C,** Overview of image segmentation pipeline of 3D-SIM data for assignment of objects (particles) for downstream measurements of features.

**D,** Outline of image segmentation pipeline with the example of CIZ1 signals, showing the output of pre- and post-filtering steps for assigning protein particles. 3D Gaussian and TopHat kernels are applied for contrast enhancement and a 3D-seeded watershed algorithm is used to obtain segmented particles. Histograms indicate the intensity profiles obtained under the dashed line during the processing steps, showing the increased resolving and separation of individual particle signals.

**E,** Pipeline for the creation of masks to include all protein signals quantified in 3D-SIM particles analyses.

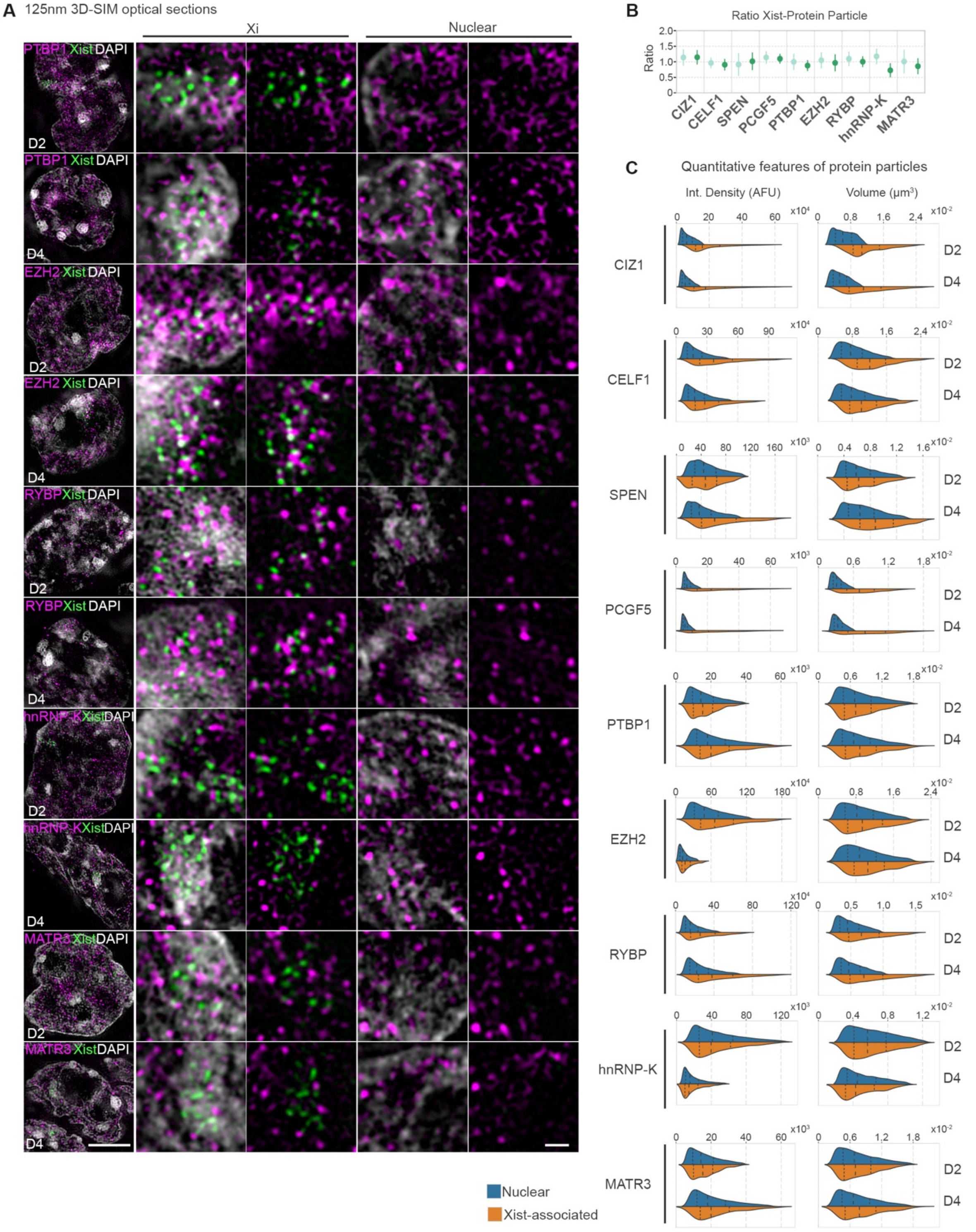


**Figure S6. Quantitative features of proteins in nuclear and Xist-associated fractions**

**A**, Left panels: 3D-SIM 125nm optical sections of cells at days 2 (D2) and 4 (D4) of differentiation showing Xist$^{MS2-GFP}$ signals (green) and immunodetected or Halo-fused proteins labelled with primary and secondary CF568-conjugated antibodies or JF549 Halo ligands (magenta) (see methods). DAPI counterstaining shown in grey. Right panels: magnifications of the Xi and nuclear regions with or without DAPI. Bar: 5µm; Magnifications: 1µm.

**B,** Point-plots showing the average ratio of the number of segmented protein particles to Xist foci within 250nm.

**C,** Violin plots of integrated densities and volumes of nuclear and Xist-associated protein particles from experiment shown in **Figures S6A** and **3E,** at days 2 and 4 of differentiation. Long-dashed black lines denote the median, short- dashed lines denote the upper/lower quartiles.

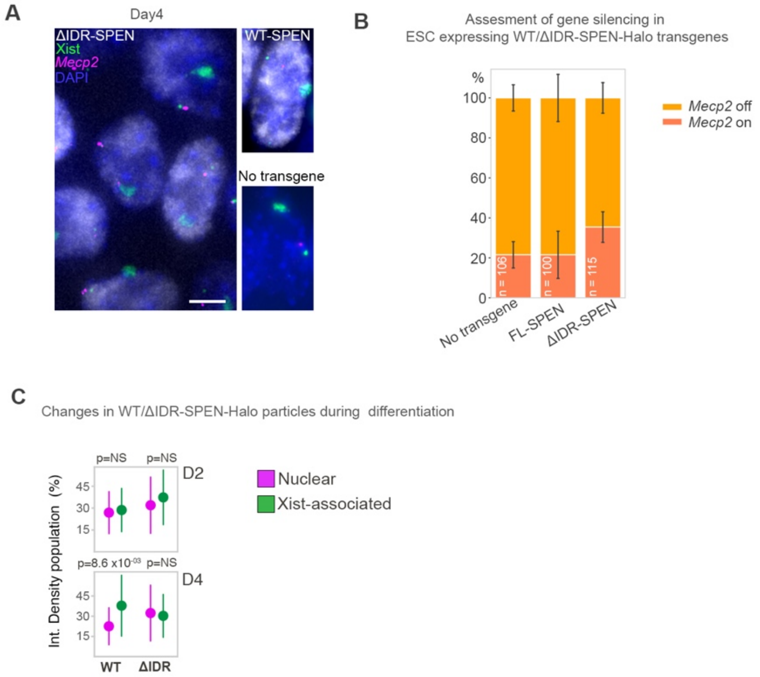


**Figure S7. Assessment of gene silencing and protein levels in differentiated cells expressing the WT or ΔIDR-SPEN transgene**

**A,** Left: RNA FISH experiment with Xist probes (green), and *Mecp2* probes (magenta) on Xist$^{MS2\text{-}GFP}$ cells expressing the ΔIDR-SPEN-Halo transgene at day 4 of differentiation. ΔIDR-SPEN-Halo signals are detected with JF549 ligand (grey) and chromatin is counterstained with DAPI (blue). Right: Same RNA FISH experiment for Xist$^{MS2\text{-}GFP}$ cells expressing the WT-SPEN-Halo transgene (top) and lacking a transgene (no transgene, bottom). Bar: 5µm.

**B,** Demonstration of effective X-linked gene silencing capacity of Xist$^{MS2\text{-}GFP}$ cells expressing ΔIDR-SPEN-Halo or WT-SPEN-Halo transgenes or cells without transgene expression.

**C,** Particle integrated density measurements, comparing data for ΔIDR-SPEN and WT-SPEN at D4 of differentiation.

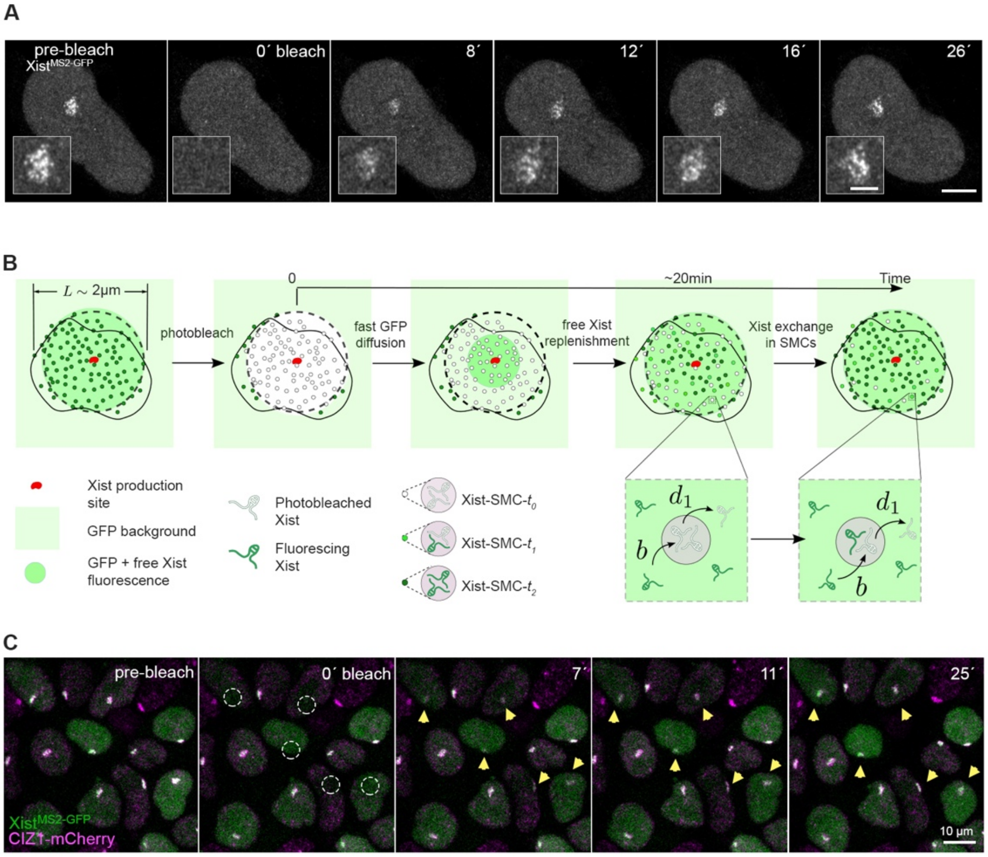


**Figure S8. Xist and CIZ1 exhibit similar kinetics**

**A,** Image sequence from FRAP experiment of $Xist^{MS2-GFP}$ at differentiation day 4, showing Airyscan optical sections. Insets show magnifications of the Xist territory. Bar: 5µm; inset: 2µm.

**B,** Schematic of the model for the Xist FRAP process. The expression and replenishment of Xist from its expression site, and of proteins from other chromosomes the nuclear fraction is assumed to be fast (free MCP-GFP replenishment is even faster; we assume is almost instantaneous just after $t$=0). The exchange of photobleached Xist with fluorescing Xist is fast in the Xi-territory outside of Xist-SMCs (free pool) and slow within Xist-SMCs. Xist-SMCs with zero, one, and two fluorescing Xist molecules are denoted Xist-SMC-0, -1, and -2, respectively. Binding of Xist to sites in the SMCs occurs at rate $b$, while dissociation occurs at rate $d_1$, which sets the timescale for FRAP recovery. The FRAP curves for Xist were fit with a single exponential. Besides the dissociation rates $d$, the percentage of fluorescence coming from freely diffusing and SMC-associated compartments, $1$-$f_1$ and $f_1$, were also inferred. See **Text S4** for modeling details.

**C,** Image sequence showing projections from FRAP experiment of $Xist^{MS2-GFP}$ cells expressing CIZ1-mCherry at D4. $Xist^{MS2-GFP}$ is shown in green and CIZ1-mCherry in magenta. Dashed circles indicate bleached Xist-territories and yellow arrows monitor recoveries of bleached regions. Bar: 10µm.

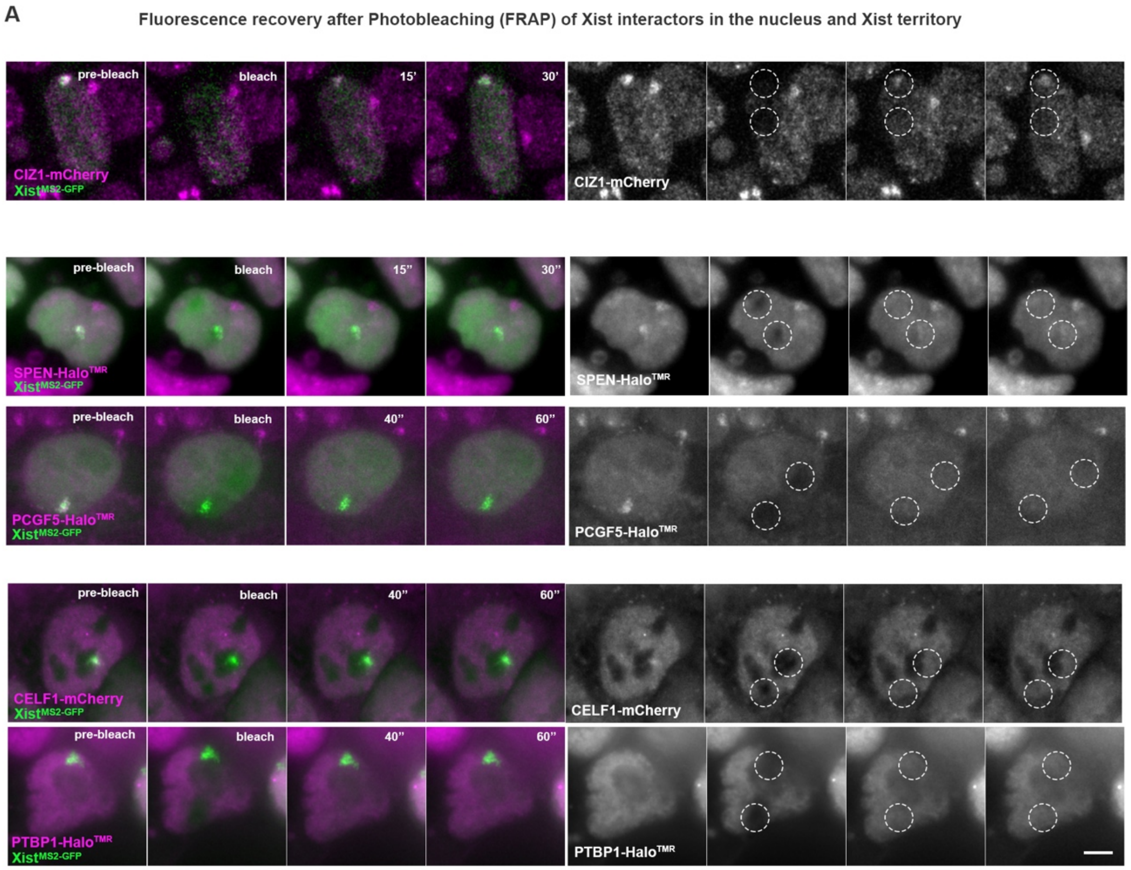


B

| | SPEN | | PCGF5 | | CELF1 | | PTBP1 | |
|---|---|---|---|---|---|---|---|---|
| | Nuclear | Xi | Nuclear | Xi | Nuclear | Xi | Nuclear | Xi |
| Fraction 1 ($f_1$) | 28 % | 23 % | 33 % | 22 % | 33 % | 18 % | 32 % | 30 % |
| Fraction 2 ($f_2$) | 10 % | 19 % | 29 % | 52 % | 31 % | 49 % | 44 % | 42 % |

**Figure S9. FRAP experiments of Xist interactors in nuclear and Xi-regions**

**A,** Time series from FRAP experiments showing bleaching (dashed circles) of nuclear or the Xi regions demarcated by Xist$^{MS2-GFP}$ signals, for CIZ1-mCherry, CELF1-mCherry, SPEN-Halo, PCGF5-Halo and PTBP1-Halo transgenes at day 4 of differentiation. Bar: 5µm.

**B,** FRAP parameters extracted from fitting for indicated proteins in the nucleus and the Xi, respectively, showing the percentage of slow ($f_1$) and fast ($f_2$) detaching fractions inferred for each protein.

A

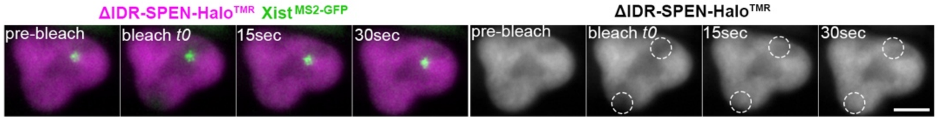


B

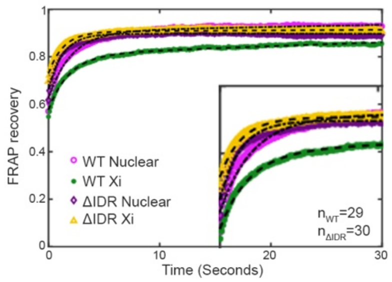


C

| ΔIDR | Lifetime | | Percentage | |
|---|---|---|---|---|
| | Nuclear | Xi | Nuclear | Xi |
| Fraction $f_1$ | 1.3sec | 1.3sec | 23% | 18% |
| Fraction $f_2$ | ∞ | 5.7min | 10% | 9% |

| WT | Lifetime | | Percentage | |
|---|---|---|---|---|
| | Nuclear | Xi | Nuclear | Xi |
| Fraction $f_1$ | 2sec | 1.9sec | 28% | 23% |
| Fraction $f_2$ | 80sec | 91.7sec | 10% | 19% |

**Figure S10. Abolishment of Xi-immobility and increased detachment rates upon deletion of the IDR domain of SPEN**

**A,** Time-sequence from FRAP experiment of $Xist^{MS2\text{-}GFP}$ cells expressing the ΔIDR-SPEN-Halo transgene at D4 of differentiation. Bar: 5µm.

**B,** FRAP recovery curves comparing WT and ΔIDR-SPEN-Halo recoveries in nuclear and Xist territories. The first 12 seconds of this graph are shown as a magnified inset.

**C,** FRAP parameters extracted from fitting for indicated proteins in the nucleus and the Xi, respectively, showing the lifetime and percentage of slow ($f_1$) and fast ($f_2$) detaching fractions inferred for each protein. Top panel are parameters inferred for ΔIDR and bottom panel for WT SPEN for comparison. Note, that the data for WT SPEN are the same shown in **Figure 3** and **Figure S9**, for comparison purposes.

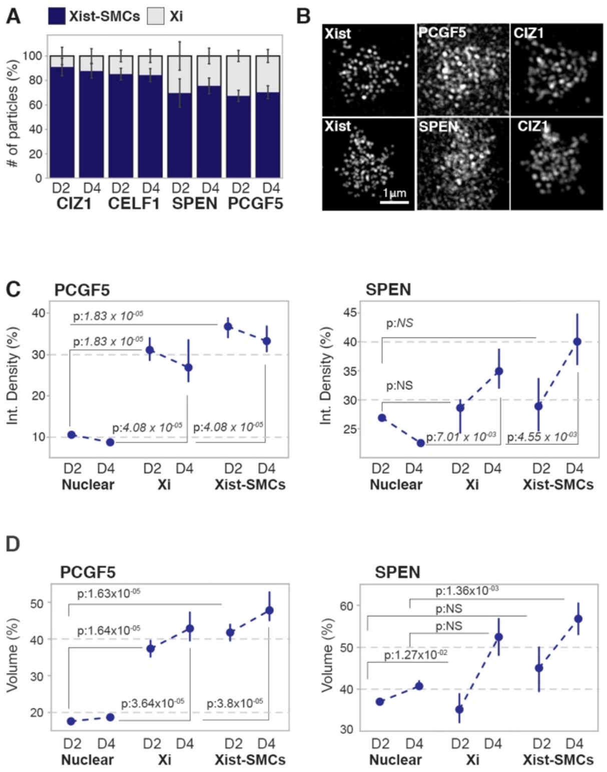


**Figure S11. Influx of proteins from Xist-SMCs regulates protein levels in the X chromosome**

**A,** Bar graphs showing the percentage of protein particles detected in the pre-Xi/Xi outside Xist-SMCs (labelled as Xi) or in Xist-SMCs, at D2 and D4. Error bars indicate standard deviation.

**B,** 3D-SIM projections showing magnifications of the Xi region for Xist, PCGF5 and CIZ1 (top) and Xist, SPEN and CIZ1 (bottom). Note the extended population of PCGF5 and SPEN in the Xi compared to Xist and CIZ1.

**C,** Point-plots showing the integrated densities of protein particles for PCGF5 (left) and SPEN (right) within the nuclear fraction, the Xi (outside Xist-SMCs) or in Xist-SMCs, at D2 and D4. Dots denote the median, bars the 95% confidence interval. The medians at days 2 and 4 are connected by dotted lines to visualize any changes. Data are normalized to the highest signal observed across the entire population. MWW *p*-values for the comparison between the nucleus and the Xi or Xist-SMCs at day 2 or 4 are given.

**D,** Same as **C** expect showing volume measurements.

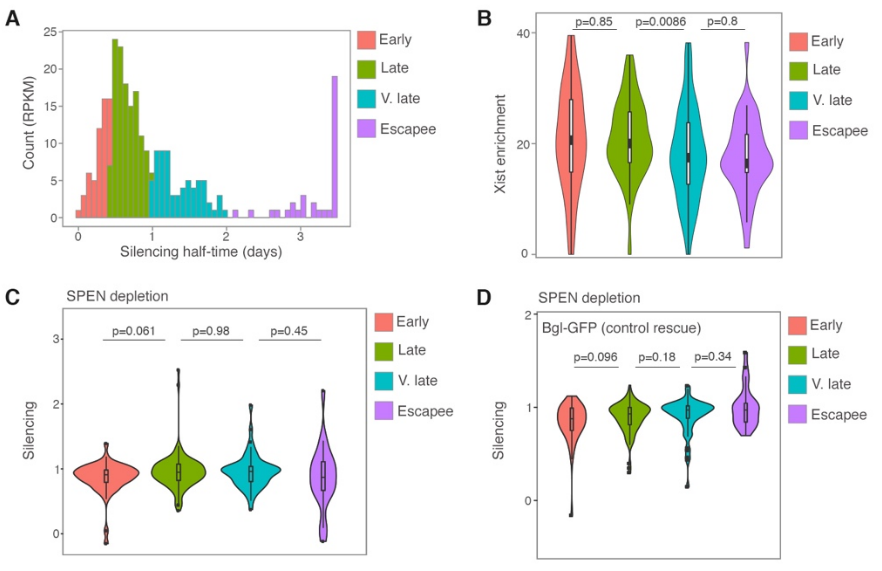


**Figure S12. SPEN-dependence of X-linked gene silencing and Xist enrichment for genes with differing silencing kinetics**

**A,** Distribution of silencing half-times of X-linked genes as determined in (*29*) and classification in early, late, and very late (v. late) silenced genes as well as XCI escapees.

**B,** Violin plots showing Xist enrichment (D2) for gene groups described in **A.** Wilcoxon *p*-value is given.

**C,** Violin plots showing silencing profiles of the gene groups defined in **A** in SPEN depleted cells. 0 indicates complete silencing and 1 complete lack of silencing. Wilcoxon *p*-value is given.

**D,** Violin plots showing the effect on gene silencing upon SPEN depletion and GFP is recruited to Xist via the BglG/SL tethering. X-linked genes are classified according to their silencing half-times during normal XCI as given in **A**. 0 indicates complete silencing and 1 complete lack of silencing by $Xist^{GFP}$ (Bgl-GFP). Wilcoxon *p*-value is given. This figure serves as control for the $Xist^{SPOC}$ (Bgl-GFP-SPOC) experiment in Figure 4B.

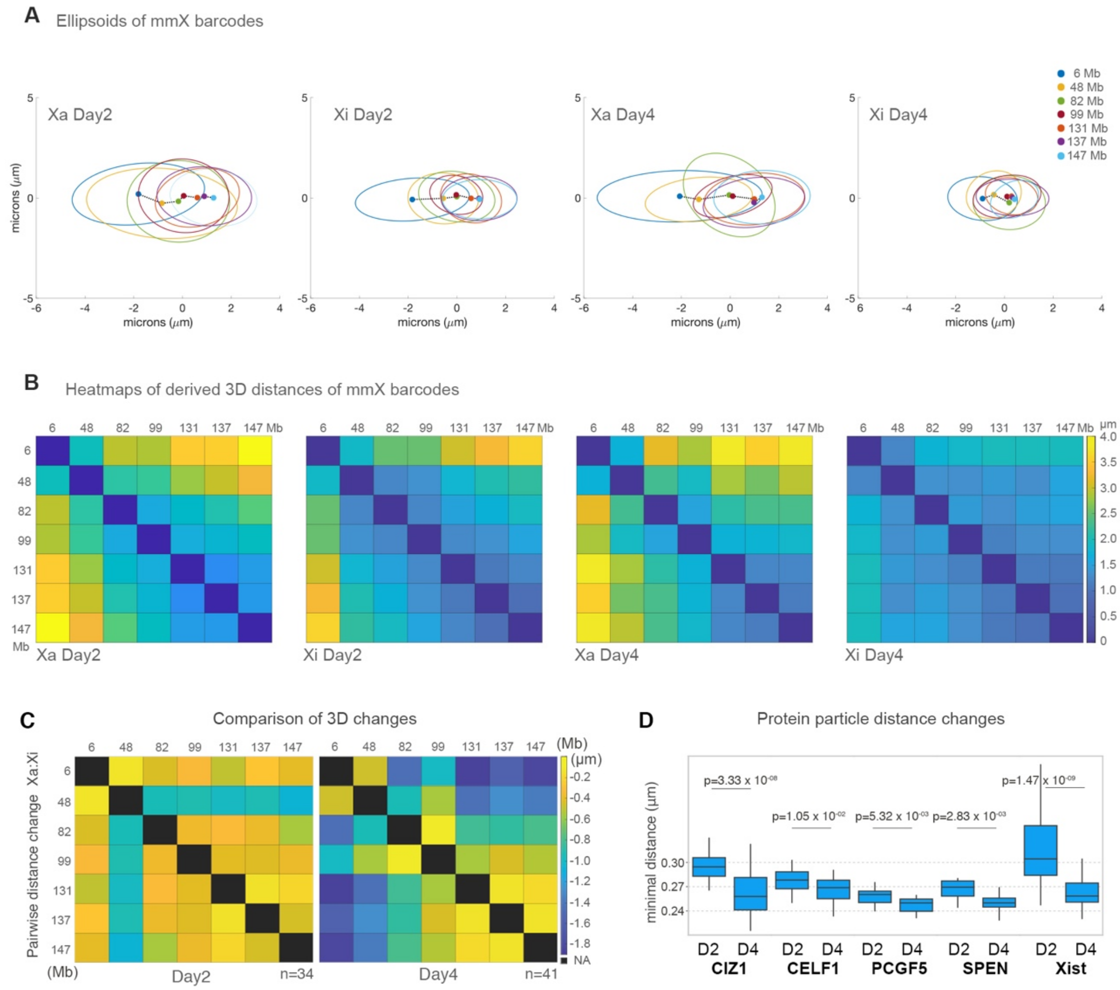


**Figure S13. Progressive compaction of the Xi and concentration of Xist-SMCs during differentiation**

**A,** Ellipsoids of 2D coordinates of X-chromosome (mmX) barcodes on the Xa and Xi at D2 and D4 of differentiation, showing a radius of 95% confidence in the allocated positions. The position along the X-chromosome for each probed location is given in megabases (Mb).

**B,** Heatmaps showing the average 3D spatial distances of genomic barcodes across the X chromosome on the Xa and Xi at D2 and D4 of differentiation.

**C,** Heatmaps as in **B,** showing changes in distances between probes on the Xa and Xi on D2 and D4.

**D,** Nearest neighbor Xist foci and protein particle distances in the Xi at D2 and D4.

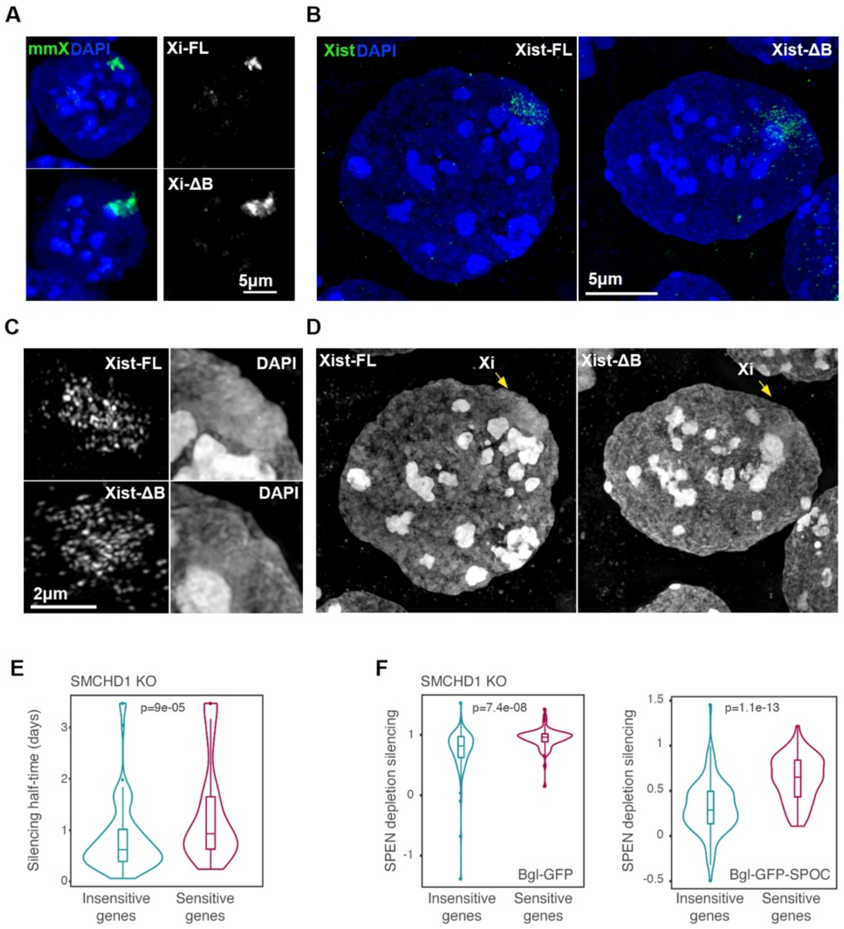


**Figure S14. Gene silencing originates at SMCs and proceeds via PRC1-mediated chromosome compaction**

**A,** Projection of DNA FISH with X-chromosome paints (mmX) in ESCs expressing tet-inducible full-length Xist (FL-Xist) or a deletion-mutant of the B-repeat (ΔB-Xist) after 18hrs of doxycycline induction. DAPI counterstaining is shown in blue.

**B,** 3D-SIM projections of RNA FISH with Xist probes (green) in WT (FL-Xist) or a deletion-mutant of the B-repeat (ΔB-Xist) in female cells at D4 of differentiation. DAPI counterstaining is shown in blue.

**C,** Magnified insets from **B** showing Xist or DAPI signals. Note the characteristic compaction of the Xi territory evident in WT cells, which is not present in ΔB-Xist cells.

**D,** As in **C** showing only the DAPI channel. The Xi regions are indicated by arrows.

**E,** Violin plots showing the silencing half-times (in days) of SCMHD1-sensitive and insensitive X-linked genes. Wilcoxon *p*-value is given.

**F,** Violin plots showing the effect on gene silencing when WT-SPEN is depleted and GFP or the SPOC-domain of SPEN is recruited to Xist via the Bgl tethering, for X-linked genes sensitive and insensitive to SCMHD1 deletion, respectively. 0 indicates complete silencing and 1 complete lack of silencing by $Xist^{GFP}$ or $Xist^{SPOC}$. Wilcoxon *p*-value is given.

| Protein pairs | number of cells | number of particles |
|---|---|---|
| **CIZ1-CELF1, D2** | 18 | 22392 |
| **CIZ1-CELF1, D4** | 17 | 8111 |
| **CIZ1-SPEN, D2** | 9 | 15992 |
| **CIZ1-SPEN, D4** | 8 | 3601 |
| **CIZ1-PCGF5, D2** | 13 | 11756 |
| **CIZ1-PCGF5, D4** | 15 | 3285 |
| **CELF1-PCGF5, D2** | 12 | 16783 |
| **CELF1-PCGF5, D4** | 11 | 15276 |

**Table S1.**
A list of the number of cells and protein particles analyzed in **Figure 3C** is given.

| Protein | number of cells | number of particles |
|---|---|---|
| **CIZ1, D2** | 47 | 22508 |
| **CIZ1, D4** | 35 | 13736 |
| **CELF1, D2** | 23 | 25581 |
| **CELF1, D4** | 31 | 37904 |
| **SPEN, D2** | 16 | 19788 |
| **SPEN, D4** | 10 | 8741 |
| **PCGF5, D2** | 12 | 12902 |
| **PCGF5, D4** | 11 | 22580 |
| **PTBP1, D2** | 11 | 21905 |
| **PTBP1, D4** | 9 | 11815 |
| **EZH2, D2** | 16 | 33851 |
| **EZH2, D4** | 13 | 12734 |
| **RYBP, D2** | 14 | 16552 |
| **RYBP, D4** | 15 | 11688 |
| **hnRNP-K, D2** | 12 | 22662 |
| **hnRNP-K, D4** | 13 | 11688 |
| **MTR3, D2** | 17 | 47314 |
| **MTR3, D4** | 18 | 89464 |
| **ΔIDR SPEN, D2** | 16 | 49283 |
| **ΔIDR SPEN, D4** | 20 | 57429 |

**Table S2.**
A list of the number of cells and protein particles analyzed in **Figures 3D-E** and **4H** is given.

| Protein | Location | Feature | D2/D4 *p*-value |
|---|---|---|---|
| **CIZ1** | Nuclear | Adjusted_IntDen | 5.98e-07 |
| **CIZ1** | Xist_Associated | Adjusted_IntDen | 1.56e-01 |
| **CIZ1** | Nuclear | Adjusted_Volume | 5.50e-08 |
| **CIZ1** | Xist_Associated | Adjusted_Volume | 3.00e-01 |
| **CELF1** | Nuclear | Adjusted_IntDen | 6.18e-03 |
| **CELF1** | Xist_Associated | Adjusted_IntDen | 1.94e-06 |
| **CELF1** | Nuclear | Adjusted_Volume | 1.64e-01 |
| **CELF1** | Xist_Associated | Adjusted_Volume | 4.01e-06 |
| **SPEN** | Nuclear | Adjusted_IntDen | 9.83e-02 |
| **SPEN** | Xist_Associated | Adjusted_IntDen | 2.12e-02 |
| **SPEN** | Nuclear | Adjusted_Volume | 1.11e-01 |
| **SPEN** | Xist_Associated | Adjusted_Volume | 3.05e-03 |
| **PCGF5** | Nuclear | Adjusted_IntDen | 2.77e-05 |
| **PCGF5** | Xist_Associated | Adjusted_IntDen | 1.70e-01 |
| **PCGF5** | Nuclear | Adjusted_Volume | 2.98e-02 |
| **PCGF5** | Xist_Associated | Adjusted_Volume | 4.38e-03 |
| **PTBP1** | Nuclear | Adjusted_IntDen | 3.12e-03 |
| **PTBP1** | Xist_Associated | Adjusted_IntDen | 7.53e-03 |
| **PTBP1** | Nuclear | Adjusted_Volume | 3.80e-01 |
| **PTBP1** | Xist_Associated | Adjusted_Volume | 2.47e-01 |
| **EZH2** | Nuclear | Adjusted_IntDen | 2.83e-06 |
| **EZH2** | Xist_Associated | Adjusted_IntDen | 2.83e-06 |
| **EZH2** | Nuclear | Adjusted_Volume | 8.18e-02 |
| **EZH2** | Xist_Associated | Adjusted_Volume | 1.10e-01 |
| **RYBP** | Nuclear | Adjusted_IntDen | 2.55e-06 |
| **RYBP** | Xist_Associated | Adjusted_IntDen | 3.13e-06 |
| **RYBP** | Nuclear | Adjusted_Volume | 5.51e-02 |
| **RYBP** | Xist_Associated | Adjusted_Volume | 1.06e-01 |
| **hnRNP-K** | Nuclear | Adjusted_IntDen | 1.25e-05 |
| **hnRNP-K** | Xist_Associated | Adjusted_IntDen | 1.25e-05 |
| **hnRNP-K** | Nuclear | Adjusted_Volume | 1.16e-05 |
| **hnRNP-K** | Xist_Associated | Adjusted_Volume | 1.52e-05 |
| **MTR3** | Nuclear | Adjusted_IntDen | 3.40e-07 |
| **MTR3** | Xist_Associated | Adjusted_IntDen | 3.49e-05 |
| **MTR3** | Nuclear | Adjusted_Volume | 3.79e-02 |
| **MTR3** | Xist_Associated | Adjusted_Volume | 1.20e-01 |

**Table S3.**

A list of the p-values derived from a Mann-Whitney Wilcoxon (MWW) test comparing the integrated density and volume of Xist-associated or nuclear fractions of Xist interactors in days 2 and 4 of differentiation is given.

| Protein | Location | MWW *p*-value |
|---|---|---|
| **CELF1** | Nuclear | 6,84e-04 |
| **CELF1** | Xi | 6,84e-04 |
| **CIZ1** | Nuclear | 2,19e-02 |
| **CIZ1** | Xi | 3,45e-04 |
| **PCGF5** | Nuclear | 5,32e-03 |
| **PCGF5** | Xi | 1,96e-02 |
| **SPEN** | Nuclear | 2,85e-01 |
| **SPEN** | Xi | 2,52e-04 |

**Table S4.**

A list of the p-values derived from a Mann-Whitney Wilcoxon (MWW) test comparing the change in density of Xi or nuclear fractions in days 2 and 4 of differentiation is given.

| Figures | Fluorescently labelled probes used in this study |
| --- | --- |
| **Fig. S1, B and C** | Xist-Atto488, Mecp2-Cy3, Atrx-Cy3 |
| **Fig. S1D** | MS2-Atto488, Xist-Cy3, Atrx-Cy5 |
| **Fig. S3A** | Xist-Atto488, Xist Intron1-Cy3 |
| **Fig. S2A-D** | Xist-Atto488 |
| **Fig. S2E** | XIST-Atto488 |
| **Fig. 4F** | Color-coded as: Green-Atto488, Yellow-Cy3, Red-Texas Red, Purple-Cy5 |
| **Fig. 4I, Fig. S13E** | mmX-Atto488 |
| **Fig. 4J, Fig. S13F-H** | Xist-Atto488 |
| **Fig. S7A** | Xist-488, Mecp2-Cy5 |

**Table S5.**
A list of probe labelling with specific fluorophores in each experiment is given.

**Movie S1.**

**Xist foci are locally confined within a radius of ~250nm**

Live-cell 3D-SIM imaging of Xist-MCP-GFP signals at day 4 of differentiation. Projections of optical stacks acquired through time are shown.

**Movie S2.**

**Xist foci are persistent through time**

Xist foci from **Movie S1** after magnification with bilinear pixel size interpolation for clarity.

**Movie S3.**

**Xist foci are locally confined at chromatin borders facing the IC space**

Live-cell 3D-SIM imaging of $Xist^{MS2\text{-}GFP}$ (green) and $H2B\text{-}Halo^{JF647}$ (magenta) at day 4 of differentiation. Projections of optical stacks acquired through time are shown.

**Movie S4.**

**Xist and CIZ1 exhibit a highly correlated motion in the Xi-territory**

Live-cell 3D-SIM imaging of $Xist^{MS2\text{-}GFP}$ (green) and $CIZ1\text{-}Halo^{JF647}$ (magenta) at day 4 of differentiation. Projections of optical stacks acquired through time are shown.
